# Lysosomal LRRC8 complex regulates lysosomal pH, morphology and systemic glucose metabolism

**DOI:** 10.1101/2024.09.22.614256

**Authors:** Ashutosh Kumar, Yonghui Zhao, Litao Xie, Rahul Chadda, Nihil Abraham, Juan Hong, Ethan Feng, John D. Tranter, David Rawnsley, Haiyan Liu, Kyla M Henry, Gretchen Meyer, Meiqin Hu, Haoxing Xu, Antentor Hinton, Chad E Grueter, E. Dale Abel, Andrew W Norris, Abhinav Diwan, Rajan Sah

**Affiliations:** Department of Internal Medicine, Cardiovascular Division, Washington University School of Medicine, St. Louis, Missouri, USA; Fraternal Order of Eagles Diabetes Research Center, Iowa City, USA; Program in Physical Therapy and Departments of Neurology, Biomedical Engineering and Orthopedic Surgery, Washington University in St. Louis, St. Louis, USA; Department of Molecular, Cellular, and Developmental Biology, University of Michigan, Ann Arbor, MI, USA; Department of Internal Medicine, Division of Cardiology, University of Iowa, Iowa City, USA; Department of Internal Medicine, Division of Endocrinology and Metabolism, Iowa City, USA; Department of Molecular Physiology and Biophysics, Vanderbilt University, Nashville, TN, 37232, USA; Department of Medicine, David Geffen School of Medicine, UCLA; Stead Family Department of Pediatrics, Endocrinology and Diabetes Division, Fraternal Order of Eagles Diabetes Research Center, University of Iowa, Iowa City, IA, USA; St. Louis VA Medical Center, St. Louis, Missouri, USA

**Keywords:** SWELL1, insulin resistance, diabetes, autophagosome, endosome

## Abstract

The lysosome integrates anabolic signalling and nutrient-sensing to regulate intracellular growth pathways. The leucine-rich repeat containing 8 (LRRC8) channel complex forms a lysosomal anion channel and regulates PI3K-AKT-mTOR signalling, skeletal muscle differentiation, growth, and systemic glucose metabolism. Here, we define the endogenous LRRC8 subunits localized to a subset of lysosomes in differentiated myotubes. We show LRRC8A regulates leucine-stimulated mTOR, lysosome size, number, pH, and expression of lysosomal proteins LAMP2, P62, LC3B, suggesting impaired autophagic flux. Mutating a LRRC8A lysosomal targeting dileucine motif sequence (LRRC8A-L706A;L707A) in myotubes recapitulates the abnormal AKT signalling and altered lysosomal morphology and pH observed in LRRC8A KO cells. *In vivo*, LRRC8A-L706A;L707A KI mice exhibit increased adiposity, impaired glucose tolerance and insulin resistance characterized by reduced skeletal muscle glucose-uptake, and impaired incorporation of glucose into glycogen. These data reveal a lysosomal LRRC8 mediated metabolic signalling function that regulates lysosomal activity, systemic glucose homeostasis and insulin-sensitivity.

## Introduction

Lysosomes are single-membrane-bound acidic organelles that contribute to recycling extra- or intracellular macromolecules, regulation of nutrient sensing and serve as signaling hubs to maintain cellular homeostasis^1–3^. Cytoplasmic cargo materials or membrane bound receptor proteins are recycled by a regulated autophagic process. The endocytosed cargo material is initially sequestered within double-membrane-bound autophagosome vesicles, which subsequently fuse with late endosomes or lysosomes to form autolysosomes. This fusion process results in the incorporation of various components, such as lysosomal membrane proteins, hydrolytic enzymes, ion channels, and transporters^4–6^. Many ion channels and transporters exist within the lysosomal membrane, which regulates lysosomal membrane potential, pH, autophagy, as well as cellular signaling and systemic metabolism. For example, a loss-of-function mutation in TRPML (mucolipin 1), primarily a Ca^2+^ channel, found in late endosomes/lysosomes, result in dysfunction of lysosomal pH regulation^7^. This leads to the accumulated autophagosomes, compromised autophagy, and ultimately the development of lysosomal storage disorders^8–10^. Similarly, gain-of-function mutations in CLC-7, a Cl-/H+ exchanger found within the lysosome, results in altered lysosome morphology and an increase in autophagosomes^11^. Cells with this mutation exhibit compromised autophagy, as they are unable to effectively degrade endocytosed cargo material and form enlarged endo-lysosome compartments^11^. TMEM206, also known as the proton-activated chloride channel (PAC)^12^ or acid-sensing osmolyte regulator^13^ (ASOR) translocates from the plasma membrane to the endosome, maintaining an optimal endosomal pH (5.5-6) by releasing Cl^-^ ions from the endosomal lumen^12^. Ablation of the PAC results concentrate intra-lysosomal Cl^-^ ions, leading to hyperacidic endosomes^12^. Furthermore, the deficiency of the PAC in the lysosomal membrane, results in overacidified lysosomes, impaired protein degradation, and the aggregation of cellular cargo materials, suggesting the requirement of optimal pH regulation in late endosomes and lysosomes for preserved physiological function^14^.

LRRC8A (leucine-rich repeat-containing protein 8A, also known as SWELL1) is an essential subunit of a heterohexameric LRRC8 channel complex consisting of LRRC8A and other LRRC8 family proteins (LRRC8B/C/D/E). The LRRC8 complex contains a transmembrane pore domain and a C-terminal 15-17 leucine rich repeat domain (LRRD), and while is predominantly considered a plasma membrane ion channel^15,16^, however; it has recently been shown to be present in lysosomes and regulates cellular osmolarity^17^. In addition, an unbiased genome-wide CRISPR screen revealed that LRRC8A null cells exhibit enlarged dysfunctional lysosomes^18^. We, and others, previously demonstrated the LRRC8 complex to be involved in the regulation of various signaling pathways, including insulin-PI3K-AKT2 signaling, adipocyte size, as well as skeletal muscle differentiation, AKT-mTOR and AMPK signaling in vitro, and metabolism and systemic glucose homeostasis in vivo^19–21^. In this study, we examine the contributions of the lysosomal LRRC8A channel complex in lysosome function, including AKT-mTOR signaling, nutrient sensing, autophagy, and systemic metabolism. We demonstrate that the LRRC8A complex is endogenously present in lysosomes and regulates leucine-mediated mTOR signaling, lysosomal morphology, pH, and autophagic flux. Mutating LRRC8A C-terminal dileucine lysosomal targeting motifs to alanine in LRRC8A-L706A; L707A knock-in (LL:AA KI) mice is sufficient to reduce lysosomal localization while preserving plasma membrane translocation. Interestingly, LL:AA skeletal muscle cells recapitulate numerous features of LRRC8A KO cells in terms of altered lysosomal morphology, pH, and expression of lysosomal markers. Moreover, LL:AA KI mice demonstrate increased adiposity, impaired glucose, and insulin tolerance, with euglycemic clamps revealing systemic insulin resistance and reduced glucose uptake in skeletal muscle.

## Results

### LRRC8 complex resides in lysosomes in native tissues

Our previous work revealed that the adipocyte LRRC8 complex regulates PI3K-AKT2 signalling and adipocyte hypertrophy, and systemic glucose homeostasis^19^. Similarly skeletal muscle LRRC8 regulates AKT-mTOR, AMPK signalling, skeletal muscle size, and systemic glucose homeostasis in vivo^20^. LRRC8 dependent mTOR signalling raised the possibility that LRRC8 may regulate lysosomal signalling ^20^. To explore this possibility biochemically, in a minimalist cell-based system, we generated wild type (WT) and LRRC8A KO HEK cell lines lentivirally transduced with epitope tagged LAMP1 (LAMP1-FLAG-HA) to perform lysosomal immunoprecipitation (Lyso-IP) using an HA antibody and blotted for LRRC8A (**Fig. 1A**). In these experiments, the cell plasma membrane is ruptured to release intracellular organelles, including endolysosomes. We then incubated the lysate with HA Ab-beads to bind the HA tagged LAMP1 and pull-down LAMP1-HA containing organelles. This revealed LRRC8A to be present in HA-immunoprecipitated lysosomes in WT HEK cells but not in LRRC8A KO HEK cells (**Fig. 1A**). To test this in native tissue, we engineered a knock-in transgenic mouse that expresses epitope tagged LAMP1 (CAG-loxP-stop-loxP-LAMP1-RFP-2xFlag-TEV-HA) upon Cre-mediated excision of a floxed stop codon (**Fig. 1B**). WT mice served as a negative control. We isolated and cultured primary skeletal muscle cells from these mice and then used adenoviral Cre to induce LAMP1-RFP-2xFlag-TEV-HA expression to allow for lysosomal immunoprecipitation. The eluted protein from LAMP1-RFP-Flag-HA expressing cells (LAMP1-RFP-Flag-HA + Cre) reveal a strong LAMP1 and HA signal (**Fig. 1C**) with no LAMP1 nor HA signal in WT negative control cells (WT + Cre). These LAMP1 and HA positive lanes are negative for ER and Golgi consistent with being intact lysosomes as opposed to other organelles. Furthermore, to confirm the membrane integrity of intact lysosomes bound with HA-beads, we treated them with mild detergent (1% Triton) and collected supernatant and bead protein fraction and immunoblotted for LAMP1 protein. The released endogenous LAMP1 protein (lower band, without HA tag) in the supernatant fraction indicates that lysosomal membranes were intact during lysosomal immunoprecipitation. Importantly, these LAMP1-HA+ organelles are positive for LRRC8A (**Fig. 1C, Left**), in addition to LRRC8B and LRRC8D (**Fig. 1C**), but not LRRC8E nor LRRC8C (**Fig. 1C**). These data reveal the lysosomal LRRC8 channel in skeletal muscle is an LRRC8A/B/D heterohexameric complex.

**Figure 1.**
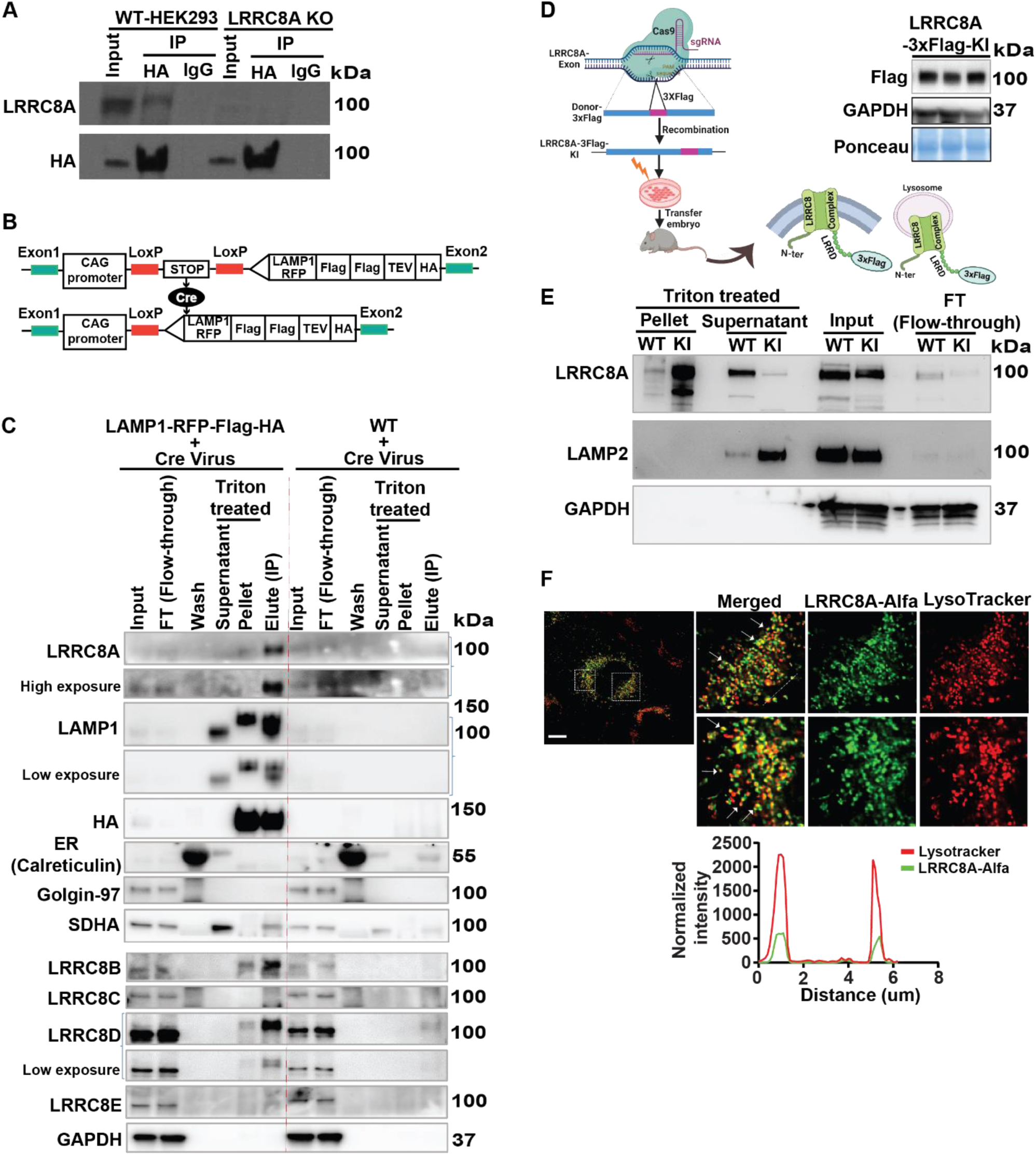
Endogenous LRRC8 proteins are present in lysosomes. **A,** Lysosomal immunoprecipitation (IP) from WT and LRRC8A KO HEK cells transiently expressing LAMP1-RFP-Flag-TEV-HA followed by LRRCA and HA western blots (WB). **B,** Schematic representation for the generation of Cre-inducible LAMP1-RFP-Flag-TEV-HA expressing mouse**. C,** Primary skeletal muscle myotubes isolated from Cre-inducible LAMP1-RFP-Flag-TEV-HA mice and C57BL/6 mice were transduced with Ad-CMV-Cre virus. After cell disruption, lyso-IP was performed with anti-HA magnetic beads to pull-down intact lysosomes and WB performed for different LRRC8 complex subunits (LRRC8A/B/C/D/E); Input (Post Nuclear Fraction), FT (Unbound protein), Wash (lysosome-bound beads washed with Wash buffer), Triton treated samples (Supernatant and pellet), Elute (Intact lysosomes bound with beads). WB of organelle-specific markers was performed to confirm the purity of isolated lysosome; ER (Calreticulin), Golgi (Golgin-97), SDAH (Mit-ComplexII), Lysosome (LAMP1), HA (epitope tag on expressed LAMP1), GAPDH (Loading control). **D,** Schematic representation of the generation of 3xFlag tagged LRRC8A knock in (KI) mouse by using CRISPR/Cas9 approach. WB with anti-flag antibody in isolated tibialis muscle of LRRC8A-3xFlag (KI) mouse. GAPDH and ponceau used as loading control (right side). **E,** Lysosomal IP performed on cardiac tissue of LRRC8A-3xFlag (KI) and WT control mice using anti-Flag magnetic beads. WB of lysosomal IP showing enriched LAMP2 protein in triton treated supernatant protein fraction of LRRC8A-3xFlag (KI) in comparison to WT lane. LRRC8A protein is enriched in the triton-treated pellet fraction, as it is bound to anti-Flag magnetic beads. GAPDH served as a loading control. **F,** Live cell confocal imaging showing colocalization of lysotracker red stained lysosomes with transiently expressed LRRC8A-ALFA (Green) in LRRC8A KO C2C12 myoblast cells. Inset image shows individual lysosome colocalizes with LRRC8A-ALFA. Scale bar: 10 µm. A line scanning intensity graph drawn between individual lysosome showing colocalization of lysosome (Red) and LRRC8A-ALFA (Green) (lower side).

As a complementary biochemical approach, we generated LRRC8A-3xFlag knock-in (KI) mice in which 3xFlag epitope was knocked into the endogenous locus of the LRRC8A C-terminus after the leucine-rich repeat domain using CRISPR/Cas9 gene-editing (**Fig. 1D**) to allow for endogenous immunoprecipitation (IP). Based on the topology of the LRRC8 channel complex, the C-terminal 3xFlag will be facing the cytoplasm, whether on the plasma membrane or within lysosomes (**Fig. 1D)** and therefore should be accessible for anti-Flag antibody mediated lysosomal IP. Endogenous LRRC8A-3xFlag protein is detectable in mouse tibialis anterior (TA) using a Flag antibody (**Fig. 1D, right**). Next, using cardiac muscle tissue from LRRC8A-3xFlag KI mice as compared to WT controls with no 3xFlag KI, we isolated intact lysosomes and performed lysosomal IP using anti-Flag antibody (**Fig. 1E**). LRRC8A protein is depleted in the flow through (FT; unbound protein) lane, indicating LRRC8A-3xFlag binding to the magnetic beads (**Fig. 1E**) with equal input loading. Application of lysis buffer to the beads released LAMP2 from LRRC8A-3xFlag bound cardiac lysosomes and much less so from WT cardiac muscle, the latter reflecting a non-specific binding of lysosome to the beads in WT samples. Subsequent bead boiling released LRRC8A-3xFlag bound to the beads confirming successful lysosomal IP from LRRC8A-3xFlag cardiac muscle (**Fig. 1E**) with minimal background in WT. As a complementary imaging approach, we generated an ALFA-epitope tagged LRRC8A construct by placing an ALFA epitope tag within the first extracellular loop yielding a versatile tool for both live cell and fixed cell imaging using high-affinity nanobodies^22^ (**Supplementary Figure 1A**). To confirm that ALFA-tagged LRRC8A forms a functional channel complex, we co-transfected LRRC8A-ALFA-IRES-EGFP and LRRC8C-P2A-mCherry in quintuple LRRC8A/B/C/D/E KO HeLa cells (5KO) and performed whole cell patch-clamp to measure hypotonically (210 mOsm) activated VRAC current. LRRC8A-ALFA;LRRC8C channels are robustly activated by hypotonic swelling and completely inhibited by DCPIB (10 uM), a selective VRAC inhibitors (**Supplementary Figure 1B**), indicating that LRRC8A-ALFA forms a functional channel with LRRC8C. Next, we transfected LRRC8A-ALFA in LRRC8A KO C2C12 myoblasts and performed immunostaining with LAMP1 protein in PFA fixed cells. Confocal microscopy reveals LRRC8A protein colocalizing with LAMP1-positives lysosomes (**Supplementary Figure 1C**). Next, we examined LRRC8A localization in live C2C12 myoblasts by transiently expressing LRRC8A-ALFA followed by fluorescence microscopy after pulse-chase application of anti-ALFA-Alexa488 conjugated nanobody and LysoTracker staining (**Fig. 1F)**. In transfected C2C12 myoblasts, ALFA-epitope tagged LRRC8A co-localizes with LysoTracker positive organelles, consistent with lysosomal LRRC8A. A line scanning intensity graph shows that LysoTracker stained (red) puncta are also positive for LRRC8A protein (green) (**Fig. 1F, lower)**. Taken together, lysosomal IP using both LAMP1-RFP-2xFlag-TEV-HA expressing mice and LRRC8A-3xFlag KI mice, and fluorescence imaging in C2C12 myoblasts expressing ALFA-tagged LRRC8A reveal LRRC8 to be present in a subset of lysosomes.

### LRRC8A regulates leucine-stimulated AKT-mTOR

As the lysosome is central to amino-acid nutrient sensing we next examined leucine-stimulated AKT and mTOR signalling in LRRC8A KO skeletal myotubes. C2C12 myotubes exhibit robust leucine-stimulated mTOR signalling with phosphorylation of p70 S6K and S6, and this is markedly suppressed in LRRC8A null cells (**Fig. 2A&B**). We next examined this in primary skeletal myotubes isolated and differentiated from LRRC8A^fl/fl^ mice and then treated with Ad-CMV-GFP or Ad-CMVCre-GFP to generate WT and LRRC8A null primary skeletal myotubes respectively (**Fig 2C&D**). Like C2C12 myotubes, WT primary skeletal myotubes exhibit clear increases in leucine-stimulated p-mTOR, p-P70 and p-S6 with no significant increases in leucine-stimulated pAKT2, and these were all markedly abrogated in LRRC8A null myotubes (**Fig 2C&D**). As our previous work revealed LRRC8A KO C2C12 cells exhibit impaired myotube differentiation^20^, we asked if the LRRC8A-dependent leucine-mediated downstream signaling defects in LRRC8A KO C2C12 cells are a result of impaired myotube differentiation, or whether this impairment is independent of the differentiation process. To test this, we first fully differentiated C2C12 cells and then knocked down (KD) LRRC8A protein by using Ad-shLRRC8A-mCherry followed by leucine stimulation (**Fig 2E&F**). Similar to LRRC8A KO C2C12 and primary skeletal myotubes, shRNA-mediated LRRC8A KD in differentiated C2C12 myotubes reveal reductions in leucine-stimulated p-S6 protein (mTOR signaling) compared to WT control cells (**Fig 2E&F**), indicating that LRRC8A-dependent leucine signaling defects persist in differentiated myotubes.

**Figure 2.**
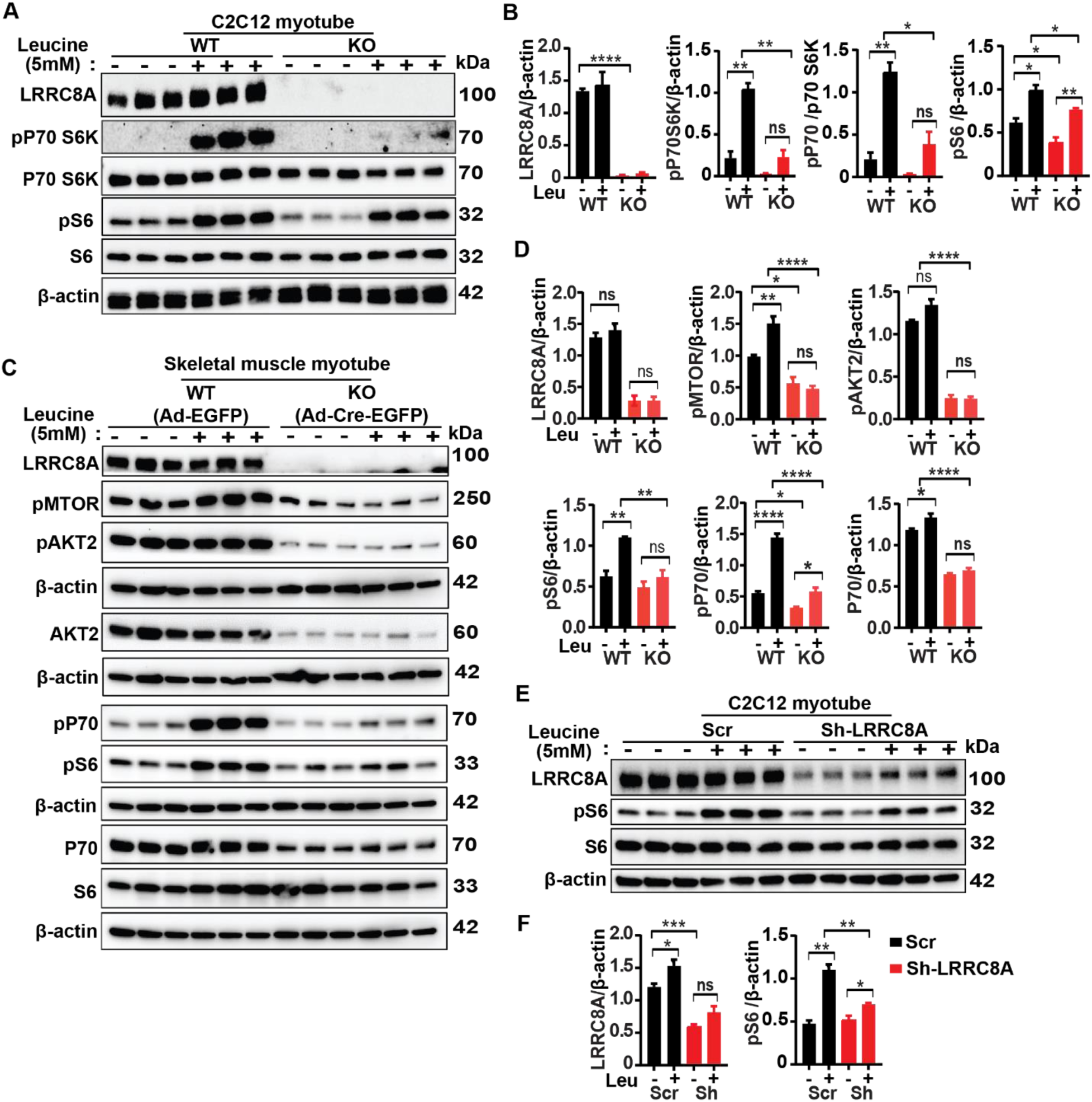
LRRC8A is required for leucine-stimulated mTOR signaling. **A,** Western blot of LRRC8A, p-P70 S6k, P70 S6K, pS6, S6, and β-actin in WT and LRRC8A KO C2C12 myotube after leucine (5 mM) stimulation for 15 minutes. **B,** Densitometry quantification of leucine-stimulated signaling WB of (**A**). **C,** WB of LRRC8A, β-actin, p-MTOR, pAKT2, AKT2, p-P70 S6k, P70 S6K, pS6, S6, protein in WT (LRRC8A^flfl^ + Ad-CMV-EGFP) and LRRC8A KO (LRRC8A^flfl^ + Ad-CMV-Cre-EGFP) primary myotube after leucine (5 mM) stimulation for 15 minutes. **D,** Densitometry quantification of leucine-stimulated signaling WB of (C). **E,** WB of LRRC8A, β-actin, and mTOR (pS6 and S6) signaling target protein in WT C2C12-sh-SCR and LRRC8A KD C2C12 myotube after leucine (5 mM) stimulation for 15 minutes. **F,** Densitometry quantification of leucine-stimulated signaling WB (E). Statistical significance between the indicated values were calculated using a two-tailed Student’s t-test. Error bars represent mean ± s.e.m. *, P < 0.05, **, P < 0.01, ***, P < 0.001, ****, P < 0.0001. n = 3, independent experiments

### LRRC8A depletion alters lysosomal size, morphology, autophagic marker protein expression and pH

To more directly examine lysosomal morphology, we performed transmission electron microscopy (TEM) imaging of WT and LRRC8A KO C2C12 myotubes (**Fig. 3A**). Lysosomes, osmiophilic structures visualized under TEM, have a 67% larger surface area and 6% reduction in circularity index in LRRC8A KO C2C12 myotubes relative to WT myotubes (**Fig. 3A**). The lysosomal circularity index is closest to 1 when lysosomes are most compact and decreases when lysosomes become irregular or aggregated. To examine this cellular phenotype in a different cell type, we performed TEM imaging in Human Umbilical Vein Endothelial Cells (HUVEC) treated with an adenoviral short-hairpin RNA (shRNA) expressing a scrambled control (Ad-shSCR) or shLRRC8A (Ad-shLRRC8A), as described previously^23^ (**Supplementary Figure 2**). Upon shRNA-mediated LRRC8A KD, HUVECs also showed enlarged autophagosomes, increased lysosomal surface area, and decreased circularity index (**Supplementary Figure 2A&B**). Next, using RNA sequencing data sets from our previously published work^19,20,23^, we compared RNA transcript of lysosomal biogenesis and autophagy associated genes in C2C12 myotubes, 3T3-F442A adipocytes and HUVECs (**Fig 3B**). These transcriptomic data from multiple different LRRC8A depleted cell types reveal markedly altered gene expression of lysosomal biogenesis (TFEB, TFEB3, LAMP1, LAMP2) and autophagy marker proteins (p62, LC-3A), implicating the LRRC8 channel complex as a regulator of lysosomal function and biogenesis. Aligned with transcriptomic data, LAMP1 and LAMP2 protein levels increase upon LRRC8A deletion in both C2C12 and primary skeletal myotubes (**Fig 3C&D**) consistent with enlarged lysosomes in LRRC8A KO C2C12 myotubes. As lysosomal proteins LAMP1 and LAMP2 regulate autophagosome fusion with lysosomes for ultimate lysosomal degradation, we investigated the autophagic marker proteins p62, LC3-I, and lipidated-LC3-II in WT and LRRC8A KO C2C12 myotubes (**Fig. 3E**). Interestingly, LRRC8A KO C2C12 myotubes show significantly increased p62 and LC3-II proteins, suggesting impaired autophagic flux despite having higher LAMPs protein. Because maintenance of optimal lysosomal pH is an important regulator of cellular function and autophagic flux, we next measured lysosomal pH in WT and LRRC8A KO C2C12 myotubes (**Fig. 3F**) and myoblasts (**Supplementary Figure 3**) using the ratiometric pH (Ex340/380) sensor Lysosensor. Both LRRC8A KO C2C12 myotubes and myoblasts have lower lysosomal pH compared to WT C2C12 (**Fig. 3G; Supplementary Figure 3B**) consistent with lysosomal acidification. As a control, treatment with the V-ATPase inhibitor Bafilomycin (Baf A1) increased pH and alkalinized lysosomes as expected (**Supplementary Figure 3A&B**). Overall, these results suggest that lysosomal function, lysosomal homeostasis, and pH are all regulated by LRRC8A channel complex.

**Figure 3.**
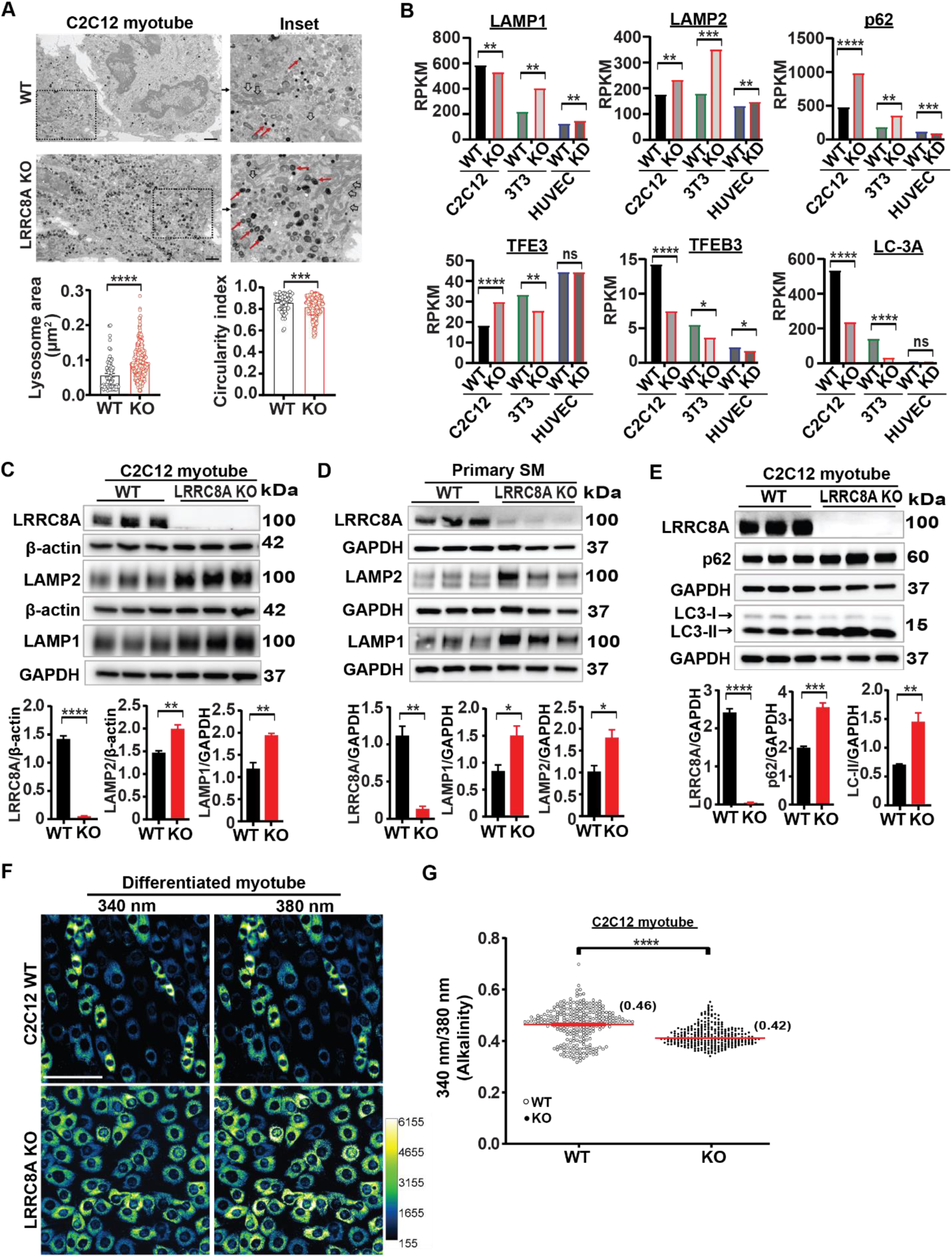
LRRC8A depletion increases lysosomal size, alters morphology and autophagic marker protein expression and decreases pH. **A,** Transmission Electron Microscopy (TEM) images of C2C12 WT and LRRC8A KO myotubes. Inset TEM image shows lysosome (red arrow) and mitochondria (Black hollow arrow). Quantification of lysosome area and circularity index (WT =73 and LRRC8A KO= 265 lysosomes). Scale bar: 2 µm. **B,** Reads Per Kilobase Million (RPKM) for autophagy and lysosome biogenesis related genes in C2C12 myotubes, 3T3-F442A adipocytes and human umbilical vein endothelial cells. **C & D,** WB of LRRC8A, LAMP1, LAMP2, β-actin and GAPDH protein in WT and LRRC8A KO C2C12 myotube (**C**) and WT (LRRC8A^flfl^ + Ad-CMV-EGFP) and LRRC8A KO (LRRC8A^flfl^ + Ad-CMV-Cre-EGFP) primary myotubes respectively (**D**) and densitometry quantification below. **E,** WB of LRRC8A and autophagy marker protein (p62, LC3-I and LC3-II) and GAPDH protein in WT and LRRC8A KO C2C12 myotubes, and densitometry quantification below. **F,** Fluorescence images of Lysosensor labeled images of WT and LRRC8A KO myotube. Scale bar: 100 µm. **G**, Ratiometric (Ex340/Ex380) intensity quantification of Lysosensor stained images from 5-6 different fields of view (n=3 independent experiment). Statistical significance between the indicated group in panel A were calculated by Mann-Whitney test. For panels B-E&G, significance between the indicated values were calculated using a two-tailed Student’s t-test. Error bars represent mean ± s.e.m. *, P < 0.05, **, P < 0.01, ***, P < 0.001, ****, P < 0.0001. n = 3, independent experiments

### Mutating the LRRC8A lysosomal targeting sequence (LRRC8A-L706A;L707A) selectively depletes LRRC8A in lysosomes

Many lysosomal membrane proteins that contain lysosomal targeting dileucine motifs as a lysosomal targeting sequence are transported to lysosomes directly via an adaptor protein 2 (AP-2)-dependent internalization process^24^. Interestingly, the LRRC8A protein C-terminus contains two consecutive dileucine motifs (LRRC8A-L706,L707), which are essential for lysosomal translocation. Mutation of these leucine residues to alanine (LRRC8A-L706A,L707A; LL:AA) is sufficient to abolish lysosomal localization while preserving plasma membrane translocation and function^17^. To explore the physiological function of lysosomal LRRC8 channels, we engineered double point mutations in the LRRC8A gene on the background of LRRC8A-3xFlag KI (KI) mice to generate LRRC8A-L706A;L707A-3xFlag-KI (LL:AA) mice (**Fig. 4A**). To confirm LL:AA mutant channels still translocate to the plasma membrane and form functional channels, we isolated skeletal muscle myoblast cells from KI and LL:AA mice and performed whole-cell patch-clamp recordings to measure hypotonically (210 mOsm) activated VRAC current. Hypotonic stimulation activated VRAC current equally in both KI and LL:AA KI myoblasts, which were both completely inhibited by DCPIB (10 uM) (**Fig. 4B&C**), indicating that LL:AA mutant forms a functional plasma membrane channel (**Fig. 4B&C**).

**Figure 4.**
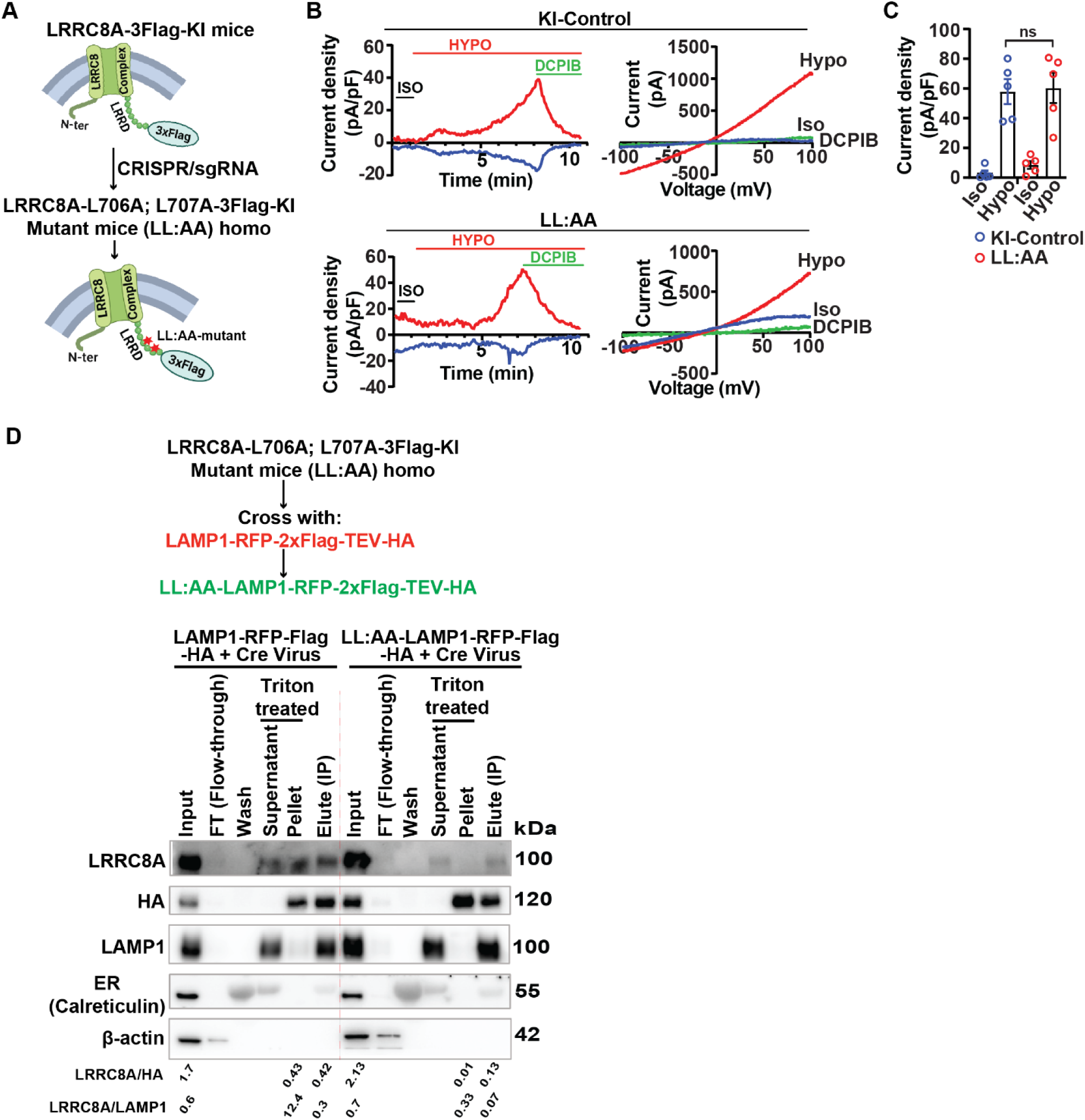
**Lysosomal targeting sequence knock-in mutation LRRC8A-L706A;L707A selectively depletes LRRC8A in lysosomes *in vivo*** **A,** CRISPR/Cas9 based approach to deplete lysosomal LRRC8 channels *in vivo* by introducing L706A;L707A double point mutations into the previously generated LRRC8A-3xFlag-KI mouse to generate LRRC8A-L706A;L707A-3xFlag KI (LL:AA) mice. **B,** Representative whole-cell patch-clamp recordings of primary myoblasts isolated from LRRC8A-3xFlag (KI-Control) and LRRC8A-L706A;L707A-3xFlag (LL:AA KI) showing current-time (left, at +100 and −100 mV) and current-voltage (right) relationships under isotonic (300 mOsm) and hypotonic (210 mOsm) conditions, followed by application of 10 uM DCPIB. Voltage protocol applied was a voltage-ramp −100 to +100 mV. **C,** Mean outward current in isotonic (300 mOsm) and hypotonic conditions (210 mOsm) recorded from KI-control (n = 5) and LL:AA KI (n = 5) myoblasts. **D,** Schematic representation for generating Cre-inducible LAMP1-RFP-Flag-TEV-HA;LRRC8A-L706A;L707A-3xFlag KI mice by crossing CAG-loxP-STOP-loxP-LAMP1-RFP-Flag-TEV-HA mice with LRRC8A-L706A;L707A-3xFlag KI mice. Primary skeletal muscle myotubes isolated from Cre-inducible LAMP1-RFP-Flag-TEV-HA mice and Cre-inducible LAMP1-RFP-Flag-TEV-HA;LRRC8A-L706A;L707A-3xFlag KI mice were each transduced with Ad-CMV-Cre and subjected to cell disruption followed by lysosomal IP with anti-HA magnetic beads to pull-down intact lysosomes. WB was performed for the LRRC8A, HA and LAMP1, ER (Calreticulin) and β-actin. The ratio of LRRC8A/HA or LRRC8A/LAMP1 protein show enrichment of LRRC8A protein in the lyso-IP lane in the WT-LAMP1 as comparison to LL:AA-LAMP1.

Next, we generated LRRC8A-LL:AA;LAMP1-RFP-HA tagged mice by crossing LRRC8A-LL:AA-3xFlag mice with transgenic LAMP1-RFP-HA mice (CAG-loxP-stop-loxP-LAMP1-RFP-2xFlag-TEV-HA) described earlier (**Fig 1B and Fig 4A**). We isolated primary skeletal muscle cells from LAMP1-RFP-HA (WT) and LRRC8A-LL:AA-LAMP1-RFP-HA mice and then induced lysosomal HA-tagged LAMP protein expression by transducing with adenoviral-Cre, followed by lysosomal IP using HA-conjugated magnetic beads. The eluted and triton treated protein from LAMP1-RFP-HA (WT) reveals the presence of LRRC8A in lysosomes (**Fig. 4D**) as observed previously (**Fig 1C and 1E**), with depletion of LRRC8A signal from LRRC8A-LL:AA-LAMP1-RFP-HA cells (**Fig. 5D**), indicating that introduction of the L706A; L707A dileucine mutation into LRRC8A-3XFlag mice selectively reduces lysosomal LRRC8 *in vivo* (**Fig. 4D**), without reducing plasma membrane LRRC8A. Indeed, densitometry analysis of LRRC8A protein signal with respect to LAMP1 (12.4 and 0.3) or HA (0.43 and 0.42) protein is 37.5 to 4.2 fold higher in LAMP1-RFP-HA (WT) lanes as comparison to LAMP1 (0.33 and 0.07) or HA (0.01 and 0.13) in LRRC8A-LL:AA mutant lanes (**Fig. 4D**). The eluted protein lane shows high LAMP1 and HA protein signal with almost no calreticulin (ER) protein signal suggesting that HA-beads bound organelles are lysosomal with minimal contamination from the ER.

**Figure 5.**
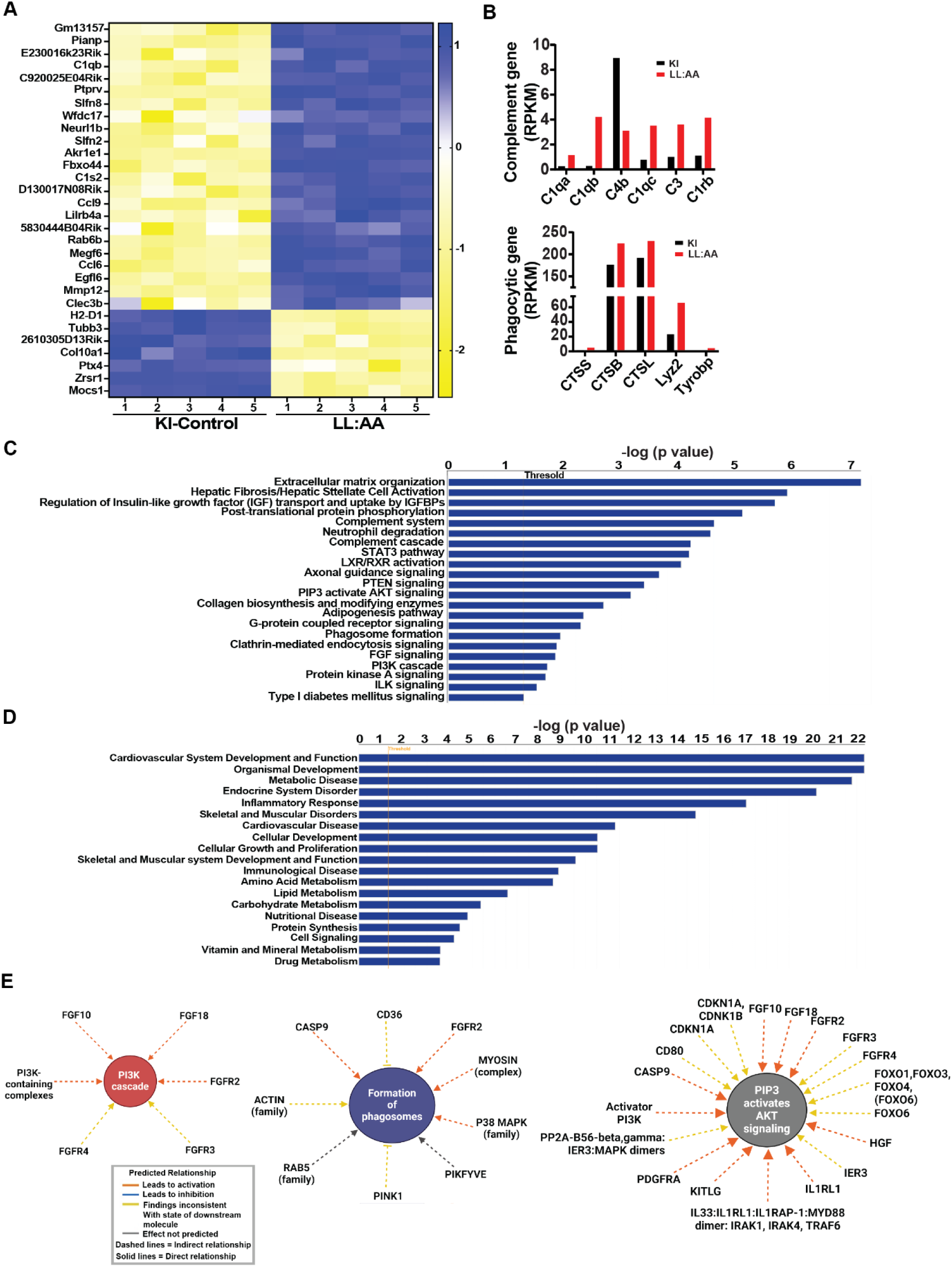
Bulk RNA transcriptome analysis of lysosomal LRRC8A depleted myotubes reveals alterations in phagosome, inflammation and metabolism associated genes. **A,** Heatmap analysis of top 30 differentially expressed genes derived from RNA transcript of LRRC8A-3xFlag KI (n=5) and LL:AA-3xFlag KI primary myotubes (n=5). **B,** Reads Per Kilobase Million for selected gene of complement cascade, lysosomal cathepsin and phagocytic marker associated genes. **C-D,** Ingenuity Pathway Analysis (IPA) of canonical pathways showing altered cellular signaling (**C**) and altered development, endocrine and metabolic pathways (**D**) in LL:AA-3xFlag KI (n=5) myotubes compared to LRRC8A3xFlag-KI (n=5). For analysis with IPA, fold change of ≥1.5, and false discovery rate <0.05 were utilized for significant differentially expressed genes. **E,** Interaction networks identified by IPA shows affected cellular signaling pathways in LL:AA-3XFlag, including PI3K cascade (left), formation of phagosomes (center), PIP3 activated AKT signaling (right). Network associated gene names and line symbol indicated by predicted legend box (lower).

### RNA sequencing of lysosomal LRRC8A depleted myotubes reveal differential expression of genes regulating inflammatory, autophagy and metabolic signalling pathways

To obtain an unbiased overview of the biological pathways impacted by selective depletion of LRRC8 channels from lysosomes we performed bulk RNA sequencing in differentiated primary myotubes isolated from KI and LL:AA mice. Interestingly, lysosomal LRRC8A depletion in primary myotubes enriched RNA transcripts of complement activation marker genes (C1q, C1qb, C4b, C1qc, C3 and C1rb) and lysosome-associated phagocytic genes such as Cathepsin (CTSS, CTSB, CTSL), Lysozyme 2 (Lyz2), and Tyrosine Motif Binding Protein (Tyrobp; DAP12) (**Fig. 5A&B, Supplementary table 1**). Hyperactivation of complement cascade associated genes and dysfunctional cathepsin enzymatic activity serve as a hallmark of a dysfunctional lysosome-autophagy axis and is linked to lysosomal storage disorders^25–29^. Furthermore, Ingenuity Pathway Analysis (IPA) of RNA transcriptome data suggest that various essential cellular signalling pathways are downregulated. These pathways include PTEN signalling (4×10^−4^), PIP3 and AKT signalling (6×10^−4^), FGF signalling (1×10^−2^), PI3K signalling (1×10^−2^) and ILK signalling (2×10^−2^) (**Fig. 5C, Supplementary table 2)**. Additionally, inflammatory or phagocytic pathways such as Complement system (2×10^−5^), Complement cascade (6×10^−5^), neutrophil degranulation (2×10^−5^), STAT3 pathway (6×10^−5^), phagosome formation (1×10^−2^) are also affected. Interestingly, these pathways are directly or indirectly associated with lysosome function (**Fig. 5C, Supplementary table 2)**. Moreover, the extracellular matrix (ECM) development pathway, which was the most highly activated canonical pathway by IPA analysis, plays a crucial role in the adhesion of muscle fibers to the ECM. This pathway is an important regulator of muscle development and homeostasis, and is regulated by lysosomal pH and the catabolic activity of lysosomes^30^. In addition, the differentially expressed genes identified by the IPA-guided network analysis, indicate various alterations in cellular signalling that are associated with metabolic disorders (**Fig. 5D & E, Supplementary Fig. 4A-D**).

### Selective depletion of lysosomal LRRC8A recapitulates defects in lysosomal morphology, pH and intracellular signalling observed in LRRC8A KO myotubes

Consistent with the pathway analysis of bulk RNA sequencing data suggesting lysosomal LRRC8 channels are important for phagosome function and biological processes that rely on phagocytic function (**Fig. 5A-E**), LL:AA myotubes phenocopy LRRC8A KO/KD myotubes and LRRC8A KD HUVEC with respect to increases in lysosome size (**Fig. 6A**), induction of LAMP1/2 expression (**Fig. 6B**), and changes in lysosomal pH (**Fig. 6C**), with no significant change in total LRRC8A protein levels (**Fig. 6B)**. These data indicate that specifically lysosomal LRRC8 localization/activity, rather than total LRRC8 activity directly regulates lysosomal function, lysosomal homeostasis, and pH in cells.

**Figure 6.**
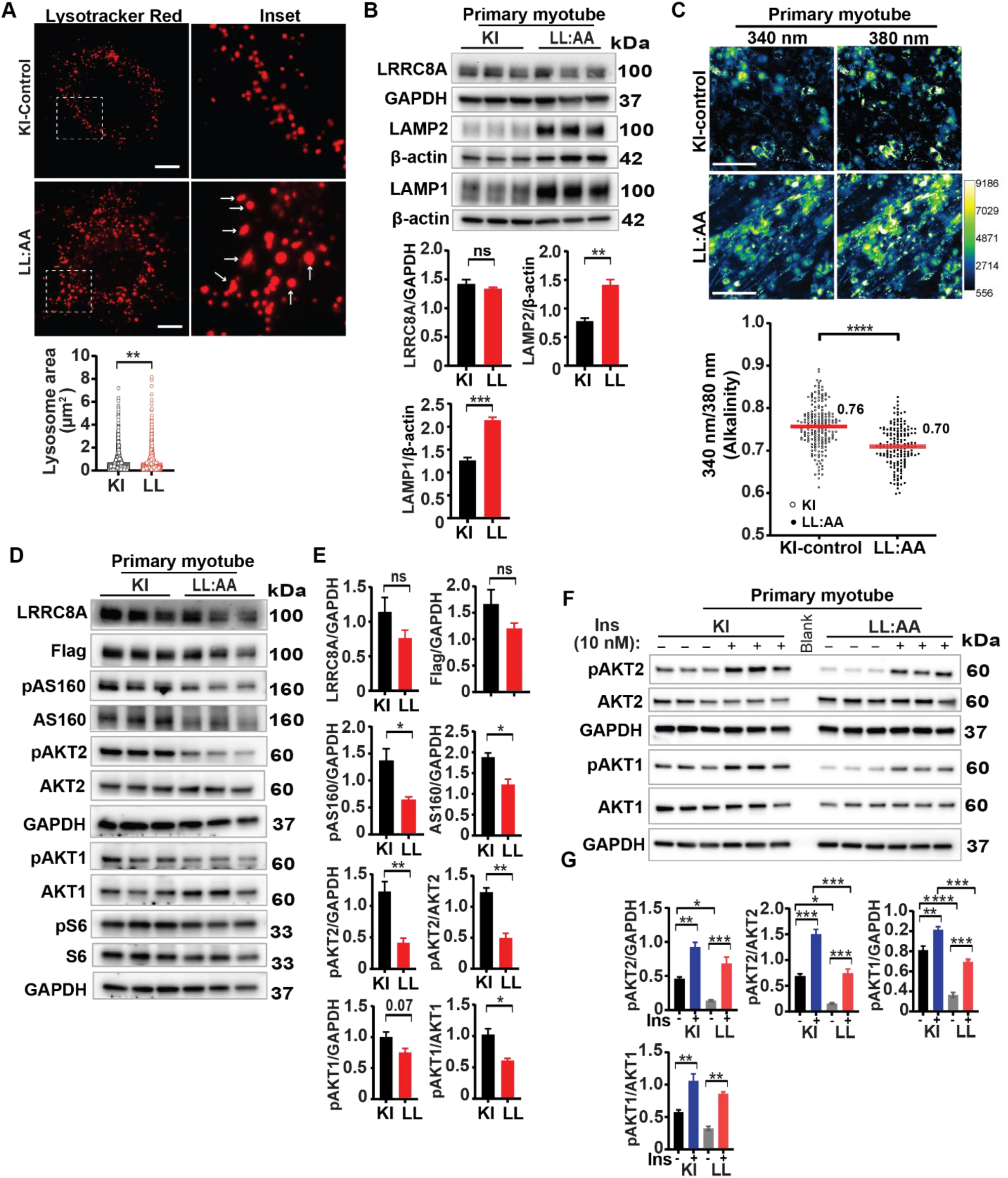
Lysosomal LRRC8A depleted myotubes phenocopy LRRC8A KO myotubes with respect to lysosomal morphology, pH and intracellular signaling. **A,** Fluorescence image of lysotracker red stained image of KI-control and lysosomal depleted LRRC8A (LL:AA) primary muscle myotubes. Scale bar: 10 um. Lysosome surface area quantified in KI-control (n= 2360 lysosomes) and LL:AA myotube (n=1813 lysosomes) images. **B,** Western blot of LRRC8A, LAMP1, LAMP2, β-actin and GAPDH protein in KI-control and LL:AA primary myotube. Densitometry quantification below. **C,** Fluorescence images of Lysosensor (Fluorescent based pH indicator) stained images of KI-control and LL:AA primary myotubes. Scale bar: 100 µm. Ratiometric (Ex340/Ex380) intensity quantification of Lysosensor stained images from 5-6 different fields of view shown below. **D,** WB of LRRC8A, Flag, pAS160, AS160, pAKT2, AKT2, pAKT1, AKT1 and GAPDH protein in KI-Control and LL:AA primary myotubes under basal conditions. **E,** Densitometry quantification of **D**. **F,** WB of pAKT2, AKT2, pAKT1, AKT1 and GAPDH protein of KI-Control and LL:AA primary myotubes stimulated with 0 and 10 nM insulin for 15 min. **G,** Densitometry quantification of **F**. Statistical significance for lysosome area was calculated by Mann-Whitney test. For all other data, statistical test for significance between the indicated values were calculated using a two-tailed Student’s t-test. Error bars represent mean ± s.e.m. *, P < 0.05, **, P < 0.01, ***, P < 0.001, ****, P < 0.0001. n = 3, independent experiments

Our previous work in the skeletal muscle cells and adipocytes suggest that LRRC8A KO cells have impaired Insulin-PI3K-AKT-mTOR signaling^19,20^. To determine if lysosomal LRRC8 channels contribute to insulin-PI3K-AKT-mTOR signaling, we isolated primary skeletal muscle cells from KI and LL:AA mice and examined the activity of these pathways under basal conditions (**Fig. 6D**). Interestingly, PI3K-AKT-pAS160 signaling is diminished in LL:AA skeletal muscle cells under basal conditions, relative to KI cells, but mTOR signaling (pS6 and S6 ribosomal protein) is unchanged, suggesting that lysosomal LRRC8A channels regulate PI3K-AKT signalling (**Fig. 6D&E**). As a complementary approach, we re-expressed WT LRRC8A or mutant LRRC8A-LL:AA in LRRC8A KO C2C12 myotubes and examined basal PI3K-AKT-mTOR signalling. Interestingly, WT LRRC8A re-expression in LRRC8A KO C2C12 myotubes fully restored PI3K-AKT-mTOR signalling while LRRC8A-LL:AA re-expression failed to restore basal PI3K-AKT-mTOR signalling (**Supplementary Fig. 5A & B**). We next examined insulin-stimulated PI3K-AKT2 signalling in KI and LL:AA primary myotubes. Consistent with the requirement of lysosomal LRRC8 channels for insulin-PI3K-AKT2 signalling, insulin-stimulated pAKT1 and pAKT2 are reduced in LL:AA myotubes compared to KI control cells (**Fig. 6F&G**). Overall, these data suggest that the lysosomal LRRC8A channel complex regulates lysosomal morphology, pH, and insulin-PI3K-AKT2 signalling in skeletal muscle cells.

### LRRC8A-LL:AA knock-in mice exhibit impaired glucose tolerance, insulin resistance and increased adiposity

In vitro experiments using LRRC8A KO C2C12 myotubes re-expressing LRRC8A-LL:AA and primary myotubes isolated from LRRC8A-LL:AA mice suggest lysosomal LRRC8 regulates lysosomal function, autophagy and insulin-PI3K-AKT2 signalling. To examine the physiological consequences of lysosomal LRRC8 depletion on systemic metabolism, we performed glucose tolerance tests (GTT) in KI and LL:AA mice raised on regular chow diet for 20-22 weeks. Glucose tolerance was significantly impaired (**Fig. 7A**) in LL:AA mice relative to KI, with no significant difference in 6h fasting glucose (**Fig. 7B**), and a trend toward increased body weight (**Fig. 7C**). This glucose intolerance was associated with impaired insulin sensitivity (**Fig. 7D**) at 22-24 weeks of age, increased fasting glucose after a 4 hour fast, and a mild, but statistically significant increase in body weight (**Fig. 7E&F**), suggesting LL:AA mice are more prone to obesity than KI mice over time. Consistent with this notion, LL:AA mice have increased body weight at 38-40 weeks of age (**Fig. 7E&F**), and body composition analysis reveals this to be due to increased adiposity, with increased fat mass, increased % fat mass, and no change in lean mass (**Fig. 7E&F**). These data indicate lysosomal LRRC8A depletion induces insulin resistance and increased adiposity with time in mice raised on a regular chow diet.

**Figure 7.**
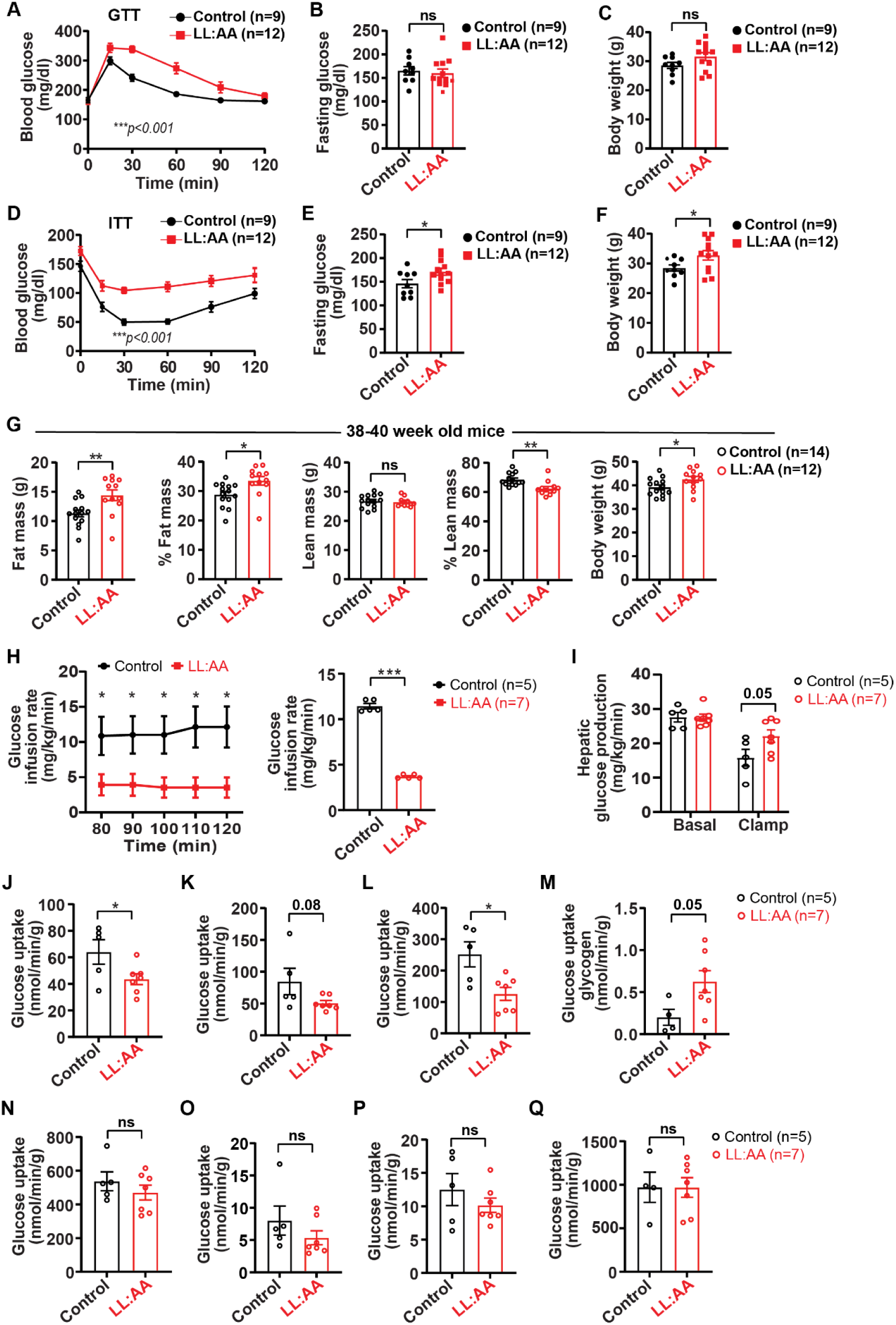
Lysosomal LRRC8A depleted mice exhibit increased adiposity, impaired glucose tolerance, insulin resistance and decreased glucose uptake. **A,** Glucose tolerance test (GTT) of KI control (n = 9) and LL:AA KI (n = 12) mice raised on chow diet for 20-22 weeks. **B,** 6 hour fasting glucose during GTT. **C,** Body weight**. D-F** Insulin tolerance test (ITT) of KI control (n = 9) and LL:AA KI (n = 12) mice raised on chow diet for 22-24 weeksn (**D**), Fasting glucose after 4 hour (**E**) and body weight (**F**). **G,** NMR measurement of absolute fat mass, % fat mass, absolute lean mass, % lean mass and body weight of KI control (n = 14) and LL:AA KI (n = 12) mice raised on chow diet for 38-40 week. **H,** Average glucose-infusion rate during euglycemic hyperinsulinemic clamps of KI control (n = 5) and LL:AA KI (n = 7) mice on chow diet for 32-36 weeks, and mean glucose-infusion rate during the entire clamp period (70-120 minutes) on right. **I,** Hepatic glucose production at baseline and during euglycemic hyperinsulinemic clamp period. **J-Q,** Glucose uptake determined from radiolabeled 2-deoxyglucose (2-DG) uptake in gastrocnemius (**J**), tibialis (**K**), soleus (**L**) muscle, liver (**M**), heart (**N**), inguinal white adipose tissue (iWAT, **O**), gonadal white adipose tissue (gWAT, **P**) and brown adipose tissue (BAT, **Q**) during traced clamp period. Data were represented as mean ± SEM. Statistical test two-way ANOVA was used for A, D and H (p-value in bottom corner of graph). Error bars represent mean ± s.e.m. *, p<0.05, **, p<0.01, ***, p<0.001. Two-tailed Student’s t-test were done for all other data. *, P < 0.05, **, P < 0.01, ***, P < 0.001, ****, P < 0.0001.

To further evaluate insulin sensitivity and determine the fate of glucose in metabolically important tissues that regulate insulin sensitivity, we performed euglycemic hyperinsulinemic clamps traced with ^3^H-glucose and 14C-deoxyglucose in in LL:AA and KI control mice raised on chow diet for 32-36 weeks. Clamp results indicate LL:AA mice require an 68% lower glucose-infusion rate (GIR) to maintain euglycemia than KI mice, indicating reduced systemic insulin sensitivity (**Fig. 7H**). The rate of glucose appearance (Ra), a measure of hepatic glucose production via gluconeogenesis or glycogenolysis, was increased 40% in LL:AA mice relative to KI mice during hyperinsulinemia, with no change at basal level, consistent with impaired suppression of hepatic gluconeogenesis, and hepatic insulin resistance (**Fig. 7I**). Next, we measured glucose uptake in individual tissues using ^14^C-deoxyglucose (2-DG). Glucose uptake was reduced in a number of skeletal muscle tissues in LL:AA mice relative to KI mice: 32% in gastrocnemius, 50% in soleus, and a non-statistically significant reduction of 40% in tibialis anterior muscle groups. There were no significant differences in glucose uptake in heart, adipose tissue (gWAT and iWAT) and brown adipose tissue (**Fig. 7J-Q**). Interestingly, incorporation of glucose into hepatic glycogen is higher in the LRRC8A-LL:AA-KI mice during the clamp period (**Fig. 7M**), despite an increase in rate of glucose appearance secondary to increased gluconeogenesis/glycogenolysis (**Fig. 7I**). Overall, these experiments reveal that LL:AA mice exhibit systemic insulin resistance primarily driven by impaired insulin sensitivity of skeletal muscle, and possibly liver, with compensatory increases in adiposity over time.

## Discussion

Lysosomes regulate various cellular processes such as signaling, nutrient sensing, autophagy, and phagosome formation. Our findings reveal that LRRC8A forms a functional lysosomal channel complex (LRRC8A/B/D), which regulates lysosomal morphology, pH, lysosome-associated membrane protein (LAMP) levels, and autophagic flux. LRRC8A depletion leads to the formation of enlarged autophagosomes. Additionally, there is evidence of impaired autophagy flux despite a lower lysosomal pH, suggesting that endocytosed material is unable to fuse with lysosomes or, even if they do fuse to form autolysosomes, the lysosomal pH is not optimal for the resident hydrolytic enzymes, resulting in accumulation of undigested material within the autophagosomes. To ensure proper enzymatic function of lysosomal hydrolases, an optimal pH range of 4.5-5.5 is essential within lysosomes. This pH balance is regulated by a variety of ion channels and pumps located on the lysosomal or endosomal membrane, including V-ATPase^31^, TMEM175^32^, TMEM206 (PAC)^12^, ClC-3-7 (acting as a 2Cl-/1H+ antiporter)^33^. It is plausible that depletion of LRRC8A protein could directly or indirectly alter the expression or canonical function of these proteins (V-ATPase, TMEM175, and ClC-3-7), resulting in deviations from optimal lysosomal pH, altered fusion of autophagosomes with lysosomes and ultimately, compromised autophagic flux. To maintain an acidic lysosomal lumen or high proton gradient, the permeation of an additional negatively charged counterion, such as Cl-, is necessary. This transport is facilitated by CLC7, which is present in the late endosome and lysosome. Notably, the presence of gain-of-function mutations in CLC7^34^ or the depletion of cellular PIP2^33^ can cause an elevation in the chloride (Cl-) gradient within the lysosomal lumen, resulting in hyperacidic lysosomes. We hypothesized that the Lyso-LRRC8A channel could act as a brake to prevent excessive acidification of the lysosome by facilitating the release of chloride ions (Cl-) from the lysosomal lumen, similar to endosomal PAC channel^12^. Additionally, an alternative hypothesis arises from a recent publication by Zhang et. al.^32^, which demonstrated lysosomal membrane proteins LAMP1 and LAMP2 directly bind to and allosterically inhibit the function of the pH-activated proton channel, TMEM175, facilitating a reduction in lysosomal pH necessary for proper lysosomal function. Based on these findings, it is possible that increased LAMP proteins in LRRC8A KO or lysosomal LRRC8 depleted LL:AA cells, could potentially bind and allosterically inhibit lysosomal TMEM175-proton channel function, thereby lowering lysosomal pH. While the mechanism underlying LAMPs protein induction in LRRC8A knockout or LL:AA cells is not yet determined, it is plausible that this upregulation may result from aggregated dysfunctional lysosomes, or possibly altered expression of genes involved in lysosomal biogenesis, such as TFEB and TFE3. Interestingly, LL:AA myotube RNAseq transcript data reveal increased lysosomal hydrolases expression (such as Cathepsin and lysozyme) as well as genes associated with the complement cascade – changes also observed in LRRC8A KO myotubes RNA seq data^20^. This suggests that abnormal protease activation in LL:AA cells could potentially trigger the activation of various complement-associated genes through site-specific cleavage. This activation may contribute to the development of autoimmune diseases and other pathological conditions. Furthermore, a recent study^35^ also showed that dysregulation of lysosomal cathepsin expression leads to the permeabilization of lysosomal membranes. This, in turn, triggers inflammasome activation, accumulation of autophagosomes, defects in autophagy, and excessive lipid accumulation in essential metabolic tissues like adipose tissue, liver, and muscle tissue^35^. These changes can contribute to obesity-related pathologies such as type 2 diabetes (T2D), fatty liver disease, and heart disease^35^. These findings are also consistent with IPA pathway analysis indicating PTEN, PI3K-AKT, IGF signaling pathways and lysosome-associated functions are altered in LL:AA myotubes.

Based on our in vitro and in vivo data, LL:AA primary myotubes phenocopy LRRC8A null myotubes with respect to defective insulin-PI3K-AKT signaling, dysfunctional lysosomes, and alteration in various metabolic pathways. These cumulative changes ultimately may lead to altered systemic metabolism and adiposity in vivo, with impaired glucose tolerance and insulin sensitivity and increased adiposity observed in LL:AA mice over time. However, there is no significant difference in the overall lean muscle mass. These results are consistent with our previous findings, in which skeletal muscle-specific LRRC8A null mice exhibit increased fat mass and insulin resistance, with no apparent difference in lean muscle mass^20^. These findings suggest that the excess circulating glucose is redistributed to other metabolic tissues, such as adipose tissue and the liver. Importantly, LL:AA mice maintain functional LRRC8 channel activity at the plasma membrane with depleted lysosomal LRRC8A protein. These mice continue to show profound abnormalities in systemic metabolism and signaling, identifying lysosomal LRRC8 as a regulator of insulin action and systemic metabolism. Numerous lines of evidence suggest that dysfunctional lysosomes, increased autophagosomes and impaired autophagy are significant contributors to the development of insulin resistance in both obesity and type 2 diabetes (T2D)^35–37^. However, the precise mechanism by which a functional lysosome regulates insulin-PI3K signaling is not yet well-established. It is plausible that the process of endocytosis of the insulin receptor and its bound ligand (insulin) and internalization into catabolic active lysosomes for degradation occurs more rapidly in LL:AA myotubes. This could be attributed to either higher expression of cathepsin enzymes^38^ or the increased acidic nature of the lysosomes, which favors more rapid dissociation of insulin from the insulin receptor^39^ in LL:AA myotubes, thereby terminating insulin signaling and contributing to insulin resistance.

In summary, our study demonstrates that the LRRC8A/B/D channel complex is present in lysosomes and regulates lysosome morphology, pH, autophagic flux, as well as PI3K-AKT signaling in vitro. Moreover, in vivo experiments highlight a novel physiological function of lysosomal LRRC8 complex in maintaining glucose homeostasis, insulin sensitivity, and adiposity under basal conditions. Further studies are needed to elucidate the mechanisms by which the lysosomal LRRC8 complex regulate lysosomal function, cellular signaling, and systemic metabolism.

| Antibodies |  |  |
| --- | --- | --- |
| Reagent / Item | Source | Information |
| Anti-AKT1 | Cell Signaling Technology | Cat #2938;<br>RRID:AB_915788 |
| Anti-AKT2 | Cell Signaling Technology | Cat #3063; RRID:<br>AB_2225186 |
| Anti-GAPDH | Cell Signaling Technology | Cat #5174; RRID:<br>AB_10622025 |
| Anti- pAS160 | Cell Signaling Technology | Cat#4288s,<br>RRID:AB_10545274 |
| Anti- AS160 | Cell Signaling Technology | Cat#2670s,<br>RRID:AB_2199375 |
| Anti- p70 S6 Kinase | Cell Signaling Technology | Cat#9202s,<br>RRID:AB_331676 |
| Anti- p-p70 S6 Kinase | Cell Signaling Technology | Cat#9205s,<br>RRID:AB_330944 |
| Anti- pS6 Ribosomal | Cell Signaling Technology | Cat#5364s,<br>RRID:AB_10694233 |
| Anti- S6 Ribosomal | Cell Signaling Technology | Cat#2317s,<br>RRID:AB_2238583 |
| Anti- flag | Sigma-Aldrich | Cat#F3165,<br>RRID:AB_259529 |
| Anti-LAMP1 | DSHB | Cat# 1d4b,<br>RRID:AB_2134500 |
| Anti-LAMP2 | DSHB | Cat# ABL-93,<br>RRID:AB_2134767 |
| Anti-SQSTM1 / p62 | Abcam | Cat# ab56416,<br>RRID:AB_945626 |
| Anti-LC3B | Novus | Cat# NB100-2220,<br>RRID:AB_10003146 |
| Anti-pAKT1<sup>Ser473</sup> | Cell Signaling Technology | Cat #9018;<br>RRID:AB_2629283 |
| Anti-pAKT2<sup>Ser474</sup> | Cell Signaling Technology | Cat #8599; RRID:<br>AB_2630347 |
| Anti-β-actin | Cell Signaling Technology | Cat #8457; RRID: AB_10950489 |
| Anti-HA | BioLegend | Cat# 901501, RRID:AB_2565006 |
| Pierce™ Anti-HA Magnetic Beads | Thermo Fisher Scientific | Cat# 88836, RRID:AB_2749815 |
| Anti-ALFA-ATTO-488 conjugated | NanaoTag biotechnologies | Cat#N1502-At488-L |
| Anti-LRRC8A | Sah Lab<sup>40</sup> (manufactured by Pacific Immunology) | N/A |
| Anti-LRRC8B | T.J. Jentsch | N/A |
| Anti-LRRC8C | Proteintech | Cat #21601-1-AP; RRID: AB_10733648 |
| Anti-LRRC8D | T.J. Jentsch | N/A |
| Anti-LRRC8E | T.J. Jentsch | N/A |
| Goat anti-Rat IgG (H+L) Secondary Antibody, HRP | Thermo Fisher Scientific | Cat# 31470, RRID:AB_228356 |
| Invitrogen Donkey anti-goat IgG (H+L)-Horseradish Peroxidase Conjugate | Thermo Fisher Scientific | Cat #A15999; RRID: AB_2534673 |
| Goat anti-rabbit IgG (H+L)-Horseradish Peroxidase Conjugate | Bio-Rad | Cat #1706515; RRID: AB_11125142 |
| **Chemicals & Kit** |  |  |
| Accutase | Corning | Cat#MT25058CI |
| Amphotericin B | Sigma-Aldrich | Cat #A2942 |
| Bafilomycin A1 | Sigma-Aldrich | Cat #B1793 |
| Bovine serum albumin | Fisher BioReagents | Cat #BP9706-100 |
| bFGF | Gibco | Cat#13256029 |
| Collagen Type II | Gibco | Cat#17101-015 |
| Clarity™ Western ECL Substrate | Bio-Rad | Cat #1705060 |
| cOmplete™ ULTRA Protease Inhibitor Cocktail | Roche Diagnostics | Cat #06538304001 |
| DAPI | Invitrogen | Cat#D1306 |
| Dispase | Roche | Cat#61206900 |
| DC™ Protein Assay Kit I | Bio-Rad | Cat #5000111 |
| DMEM/F12 | Gibco | Cat#11330-032 |
| DMEM | Gibco | Cat#11965 |
| DNase digestion kit | Thermo Fisher Scientific | Cat#12185010 |
| HumulinR | Novolin | Cat#NDC-0169-1833-11 |
| Insulin | Cell Applications | Cat#128-100 |
| Insulin-Transferrin-Selenium (ITS -G) | Thermo Fisher Scientific | Cat#41400-045 |
| 4X Laemmli Sample Buffer | Bio-Rad | Cat #1610747 |
| Lipofectamine 2000 | Thermo Fisher Scientific | Cat#11668-019 |
| LysoView 550 | Biotium | Cat#70083 |
| LysoSensor Yellow/Blue DND-160 | Thermo Fisher Scientific | Cat#7545 |
| Matrigel | Corning | Cat#356231 |
| Penicillin / streptomycin | Thermo Fisher Scientific | Cat #15140122 |
| PhosSTOP™ | Roche Diagnostics | Cat #4906845001 |
| RNA isolation (PureLink™ RNA mini kit) | Invitrogen | Cat #12183018A |
| Tris/glycine buffer | Bio-Rad | Cat #1610771 |
| Tris/glycine/SDS running buffer | Bio-Rad | Cat #1610772 |
| μMACS™ and MultiMACS™ DYKDDDDK Isolation Kits | Miltenyi Biotech | Cat#130-101-591 |
| **Cell Lines** |  |  |
| C2C12 Cell line | ATCC | CRL-1772,<br>RRID:CVCL_0188 |
| C2C12-LRRC8A KO | Sah Lab<sup>20</sup> | NA |
| HEK293 | ATCC | CRL-1573,<br>RRID:CVCL_0045 |
| HEK-LRRC8A KO | Sah Lab<sup>41</sup> | N/A |
| HeLa quintuple KO | Zhaozhu Qiu, John Hopkins, Baltimore, MD, USA | N/A |
| **Experimental Models: Mouse/Strains** |  |  |
| Mouse: C57BL/6N | Charles River Laboratories | Cat #027 |
| Mouse: *Lrrc8a*<sup>fl/fl</sup> (*Swell1*<sup>fl/fl</sup>) | Sah Lab<sup>19</sup> Washington University, Saint Louis | N/A |
| Mouse: LRRC8A-3xFlag-KI | Sah Lab, Washington University, Saint Louis | N/A |
| Mouse: LRRC8A-L706A;L707A-3xFlag-KI | Sah Lab, Washington University, Saint Louis | N/A |
| Mouse: LAMP1-RFP-1xFlag-TEV-HA | Diwan Lab, Washington University, Saint Louis | N/A |
| **Recombinant DNA and plasmids** |  |  |
| Ad5-CMV-Cre-eGFP | University of Iowa viral vector core facility | N/A |
| Ad5-CMV-eGFP | University of Iowa viral vector core facility | N/A |
| Ad5-U6-scramble-mCherry | University of Iowa viral vector core facility | N/A |
| Ad5-mCherry-U6-shLRRC8A | University of Iowa viral vector core facility | N/A |
| Ad5-mPGK-mLrrc8a-(L706A)(L707A)-3xFlag | Vector Biolabs | N/A |
| Ad5-mPGK-mLrrc8a-3xFlag | Vector Biolabs | N/A |
| Ad(RGD)-CMV-iCre | Vector Biolabs | N/A |
| LRRC8C-P2A-mCherry | Vector Biolabs | N/A |
| pLV-CMV-rLAMP-RFP-PGK-puro | Diwan Lab, Washington University, Saint Louis | N/A |
| LRRC8A-ALFA-IRES-EGFP | Brohawn lab, University of California Berkeley | N/A |
| LRRC8A-ALFA | Brohawn lab, University of California Berkeley | N/A |

| Software |  |  |
| --- | --- | --- |
| BioRender | https://www.biorender.com/ | N/A |
| Fiji | https://imagej.net/ – Schindelin *et al.*<sup>42</sup> | N/A |
| Prism | GraphPad Software | Version 10.2.0 |
| pClamp | Molecular Devices | Version 10.4 |

## Materials and methods

### Animals

The Institutional Animal Care and Use Committee of Washington University in St. Louis approved all experimental procedures involving mice. All mice were housed in temperature, humidity, and light-controlled room and allowed free access to water and food. Study mice were gender/age matched, and both male and female mice were used. C57BL/6N mice were obtained from Charles River Laboratories (Wilmington, MA, USA). All mice were fed ad libitum with regular chow (RC; NIH31 irradiated, #7913) diet.

The generation of *LRRC8A*^*fl/fl*^ conditional mouse was described previously^19^. LRRC8A-3xFlag-KI (KI) mice were generated by CRISPR/Cas9 gene-editing where a 3xFlag epitope was placed at the C-terminus of the *lrrc8a* gene at the Mouse Genetics Core (MGC facility, Washington University, St. Louis. LRRC8A-L706A;L707A-3xFlag-KI (LL:AA) mice were generated by CRISPR/Cas9 gene-editing where L706;L707 dileucines were mutated to alanines on the background of the previously generated LRRC8A-3xFlag-KI mouse to generate mice with LRRC8A-L706A;L707A-3xFlag knock-in mutations. LoxP-STOP-LoxP-LAMP1-RFP-1xFlag-TEV-HA (LAMP1-RFP-Flag-HA)^43^ conditional mice were generated in the laboratory of Dr. Abhinav Diwan, Washington University, Saint Louis. Furthermore, LRRC8A-L706A;L707A-3xFlag-KI (LL:AA) mice were crossed with LoxP-STOP-LoxP-LAMP1-RFP-1xFlag-TEV-HA(LAMP1-RFP-Flag-HA) to generate LRRC8A-L706A;L707A-3xFlag-KI;LoxP-STOP-LoxP-LAMP1-RFP-1xFlag-TEV-HA (LL:AA-LAMP1-RFP-Flag-HA) mice.

### Adenovirus/plasmids

Adenovirus type 5 with Ad5-CMV-eGFP (3×10^10^ PFU/ml), Ad5-CMV-Cre-eGFP (8×10^10^ PFU/ml), Ad5-U6-scramble-mCherry (sh-SCR) (9×10^10^ PFU/ml), Ad5-mCherry-U6-shLRRC8A (sh LRRC8A) (1×10^11^ PFU/ml) were obtained from the University of Iowa Viral Vector Core. Ad5-mPGK-mLrrc8a-(L706A)(L707A)-3xFlag (LL:AA) (1×10^10^ PFU/ml) and Ad5-mPGK-mLrrc8a-3xFlag (1×10^10^ PFU/ml), Ad(RGD)-CMV-iCre ((1×10^10^ PFU/ml) viruses were obtained from Vector Biolabs. LRRC8C-P2A-mCherry plasmid was obtained from Vector Biolab. LRRC8A-ALFA-IRES-EGFP and LRRC8A-ALFA plasmids were generated by Dr. Stephen G. Brohawn lab, University of California Berkeley, Berkeley, CA USA.

### Electrophysiology

Patch-clamp recordings were performed in the whole-cell configuration at room temperature, as described previously^19,20,40^. Briefly, currents were recorded using either an Axopatch 200B amplifier or a MultiClamp 700B amplifier (Molecular Devices), both paired with a Digidata 1550 digitizer and data were acquired by pClamp 10.4 software. The extracellular solution for hypotonic stimulation contained (in mM): 90 mM NaCl, 2 mM CsCl, 1 mM MgCl_2_, 1 mM CaCl_2_, 10 mM HEPES, 10 mM mannitol, pH 7.4 with NaOH (210 mOsm/kg). The isotonic extracellular solution had the same composition as above, except it contained 110 mM mannitol instead of 10 mM, resulting in an osmolarity of 300 mOsm/kg. The intracellular solution contained (in mM): 120 L-aspartic acid, 20 CsCl, 1 MgCl_2_, 5 EGTA, 10 HEPES, 5 MgATP, 120 CsOH, 0.1 GTP, pH 7.2 with CsOH and had an osmolarity of 280–290 mOsm/kg. The osmolarity was checked by a vapor pressure osmometer 5500 (Wescor). Currents were filtered at 10 kHz and sampled at 100 μs interval. The patch pipettes were pulled from borosilicate glass capillary tubes (WPI) using a P-87 micropipette puller (Sutter Instruments). The pipette resistance was ∼2-5 MΩ when the patch pipette was filled with intracellular solution. The holding potential was 0 mV. Voltage ramps from −100 to +100 mV (at 0.4 mV/ms) were applied every 4 s. Clampfit 10 (Molecular Devices) was used for data analysis. Currents were normalized by cell capacitance to calculate current densities.

### Primary Muscle Satellite Cell Isolation

Satellite cell isolation and differentiation were performed as described previously^20^. Briefly, gastrocnemius, tibialis, soleus and quadriceps muscles were excised from mice (8-10 weeks old) and washed twice with cold 1XPBS supplemented with 1% penicillin-streptomycin (#15140-122, Gibco) and amphotericin B (300 ul/100ml) (#30-003-CF, Corning, Amphotericin B). The excised muscle tissue was incubated in DMEM-F12 media supplemented with collagenase II (2 mg/ml) (#17101-015, Gibco) 1% penicillin-streptomycin and amphotericin B (300 ul/100ml) at 37°C water bath for 15 minutes, and subsequently incubated in a shaker (220 rpm) for 90 minutes at 37°C. Tissue was washed with warm 1XPBS supplemented with 1% penicillin-streptomycin and fungizone (300 ul/100ml) and incubated again with DMEM-F12 media supplemented with collagenase II (1 mg/ml), dispase (0.5 mg/ml) (#61206900, Roche) 1% penicillin-streptomycin and fungizone (300ul/100ml) in a shaker for 30 minutes at 37°C. Subsequently, the tissue was minced and passed through a cell strainer (100 µm), and after centrifugation; satellite cells were plated on Matrigel-coated dishes. The isolated satellite cells were stimulated to differentiate into myoblasts in DMEM-F12, 20% fetal bovine serum (FBS), 40 ng/ml basic fibroblast growth factor (bfgf, Gibco #13256029), 1X non-essential amino acids, 0.14 mM β-mercaptoethanol, 1X penicillin/streptomycin, and Amphotericin B. Myoblasts were maintained with 10 ng/ml bfgf and then differentiated in DMEM-F12, 2% FBS, 1X insulin–transferrin–selenium, when 70-80% confluency was reached.

### Cell culture and signaling studies

WT C2C12 and LRRC8A KO C2C12 cell line were cultured at 37°C, 5% CO_2_ Dulbecco’s modified Eagle’s medium (DMEM; GIBCO) supplemented with 10% fetal bovine serum (FBS; Atlanta Bio selected) and antibiotics 1% penicillin-streptomycin. Cells were grown to 80% confluency and then transferred to differentiation media DMEM supplemented with antibiotics and 2% horse serum (#16050122, Gibco) to induce differentiation. The differentiation medium was changed every two days. Cells were allowed to differentiate into myotubes for 7-8 days. For leucine stimulation, differentiated C2C12 myotubes were incubated with Krebs-Ringer Bicarbonate HEPES (KRBH) buffer [129 mM NaCl, 5 mM NaHCO3, 4.8 mM KCl, 1.2 mM KH_2_PO_4_, 2.5 mM CaCl2, 2.4 mM MgSO4, 10 mM HEPES, 30 mM mannitol, 0.1% BSA, pH 7.4] without glucose for 3 hours. Subsequently cells were stimulated either with glucose alone [129 mM NaCl, 5 mM NaHCO_3_, 4.8 mM KCl, 1.2 mM KH_2_PO_4_, 2.5 mM CaCl_2_, 2.4 mM MgSO_4_, 10 mM HEPES, 30 mM mannitol, 0.1% BSA, pH 7.4] without glucose for 3 hours. Subsequently cells were stimulated either with glucose alone [129 mM NaCl, 5 mM NaHCO3, 4.8 mM KCl, 1.2 mM KH_2_PO_4_, 2.5 mM CaCl_2_, 2.4 mM MgSO_4_, 10 mM HEPES, 24.5 mM mannitol, 5.5 mM Glucose, 0.1% BSA, pH 7.4] or Glucose + Leucine [129 mM NaCl, 5 mM NaHCO_3_, 4.8 mM KCl, 1.2 mM KH_2_PO_4_, 2.5 mM CaCl_2_, 2.4 mM MgSO_4_, 10 mM HEPES, 19.5 mM mannitol, 5.5 mM Glucose, 5 mM Leucine, 0.1% BSA, pH 7.4] for 15 minutes. The osmolarity of the Krebs buffer was maintained at 300 mOsm by changing mannitol concentration. To perform leucine stimulation in LRRC8A KO primary muscle cells, satellite cells were isolated from LRRC8A^fl/fl^ mice and transduced with Ad5-CMV-eGFP or Ad5-CMV-Cre-eGFP (MOI 50-60) for 48 hours in differentiation media. After 3 days of differentiation, leucine stimulation was performed (as described above). To examine leucine stimulation in LRRC8A KD of post-differentiated WT C2C12 cells, we first differentiated WT C2C12 cell in differentiation for 6 days, subsequently transduced with Ad5-U6-scramble-mCherry (sh-SCR) or Ad5-mCherry-U6-shLRRC8A(sh LRRC8A) (KD) (MOI 50-60) for 48 hours, allowed them to differentiate 1 additional day before using them for leucine stimulation (as described above).

For insulin stimulation, differentiated primary myotubes were incubated in serum free media (DMEM-F12, 1% PS) for 3 hours and stimulated with 0 and 10 nM insulin (#128-100, Cell Applications) in differentiation media (DMEM-F12, 2% FBS, 1X insulin–transferrin–selenium) for 15 minutes. To perform intracellular signaling in C2C12 WT and LRRC8A KO C2C12 myotube, we re-expressed LRRC8A-3xFlag (Ad5-mPGK-mLrrc8a-3xFlag) and LRRC8A-L706A; L707A-3xFlag (Ad5-mPGK-mLrrc8a-(L706A)(L707A)-3xFlag) by transduction (MOI 80-100) in LRRC8A KO C2C12 cells in differentiation media (DMEM, 2% horse serum and 1% penicillin-streptomycin) for 48 hours and allowed them to further differentiate in differentiation media for 7-8 days. Differentiated myotube were harvested in RIPA buffer at basal condition for further signaling studies. To perform basal cellular signaling in primary skeletal muscle cells, LRRC8A-3xFlag-KI and LRRC8A-L706A; L707A-3xFlag KI cells were differentiated in differentiation media (DMEM-F12, 2% FBS, 1X insulin–transferrin–selenium) for 3 days and myotubes were harvested in RIPA buffer. WT C2C12 and LRRC8A KO C2C12 cell lines were cultured at 37°C, 5% CO_2_ Dulbecco’s modified Eagle’s medium (DMEM; GIBCO) supplemented with 10% fetal bovine serum (FBS; Atlanta Bio selected) and antibiotics 1% penicillin-streptomycin. Cells were grown to 80% confluency and then transferred to differentiation media DMEM supplemented with antibiotics and 2% horse serum (#16050122, Gibco) to induce differentiation. The differentiation medium was changed every two days. Cells were allowed to differentiate into myotubes for 7-8 days. For leucine stimulation, differentiated C2C12 myotubes were incubated with Krebs-Ringer Bicarbonate HEPES (KRBH) buffer [129 mM NaCl, 5 mM NaHCO_3_, 4.8 mM KCl, 1.2 mM KH_2_PO_4_, 2.5 mM CaCl_2_, 2.4 mM MgSO_4_, 10 mM HEPES, 30 mM mannitol, 0.1% BSA, pH 7.4] without glucose for 3 hours. Subsequently cells were stimulated either with glucose alone [129 mM NaCl, 5 mM NaHCO3, 4.8 mM KCl, 1.2 mM KH_2_PO_4_, 2.5 mM CaCl_2_, 2.4 mM MgSO_4_, 10 mM HEPES, 24.5 mM mannitol, 5.5 mM Glucose, 0.1% BSA, pH 7.4] or Glucose + Leucine [129 mM NaCl, 5 mM NaHCO_3_, 4.8 mM KCl, 1.2 mM KH_2_PO_4_, 2.5 mM CaCl_2_, 2.4 mM MgSO_4_, 10 mM HEPES, 19.5 mM mannitol, 5.5 mM Glucose, 5 mM Leucine, 0.1% BSA, pH 7.4] for 15 minutes. The osmolarity of the Krebs buffer was maintained at 300 mOsm by changing mannitol concentration. To perform leucine stimulation in LRRC8A KO primary muscle cells, satellite cells were isolated from LRRC8A^fl/fl^ mice and transduced with Ad5-CMV-eGFP or Ad5-CMV-Cre-eGFP (MOI 50-60) for 48 hours in differentiation media. After 3 days of differentiation, leucine stimulation was performed (as described above). To examine leucine stimulation in LRRC8A KD of post-differentiated WT C2C12 cells, we first differentiated WT C2C12 cell in differentiation media for 6 days, subsequently transduced them with Ad5-U6-scramble-mCherry (sh-SCR) or Ad5-mCherry-U6-shLRRC8A(sh LRRC8A) (KD) (MOI 50-60) for 48 hours, allowed them to differentiate for 1 additional day before using them for leucine stimulation (as described above).

For insulin stimulation, differentiated primary myotubes were incubated in serum free media (DMEM-F12, 1% PS) for 3 hours and stimulated with 0 and 10 nM insulin (#128-100, Cell Applications) in differentiation media (DMEM-F12, 2% FBS, 1X insulin–transferrin–selenium) for 15 minutes. To perform intracellular signaling in C2C12 WT and LRRC8A KO C2C12 myotubes, we re-expressed LRRC8A-3xFlag (Ad5-mPGK-mLrrc8a-3xFlag) and LRRC8A-L706A; L707A-3xFlag (Ad5-mPGK-mLrrc8a-(L706A)(L707A)-3xFlag) by transduction (MOI 80-100) in LRRC8A KO C2C12 cells in differentiation media (DMEM, 2% horse serum and 1% penicillin-streptomycin) for 48 hours and allowed them to further differentiate in differentiation media for 7-8 days. Differentiated myotubes were harvested in RIPA buffer under basal conditions for further signaling studies. To perform basal cellular signaling in primary skeletal muscle cells, LRRC8A-3xFlag-KI and LRRC8A-L706A; L707A-3xFlag KI cells were differentiated in differentiation media (DMEM-F12, 2% FBS, 1X insulin–transferrin–selenium) for 3 days and myotubes were harvested in RIPA buffer.

### Immunofluorescence and Lysotracker Imaging

LRRC8A KO cells were cultured on coverslip and transfected with LRRC8A-ALFA plasmid by using lipofectamine 2000 (#11668-019, Thermo Fisher Scientific) as per manufacturer instructions. After 48 hours of transfection cells were fixed with 2% PFA (#J19943-K2, Thermo Scientific) for 15 minutes in an incubator. Subsequently, fixed cells were washed 3 times with 1X PBS and permeabilized with 0.2% Tritionx100 (0.2% Triton in 1X PBS) for 10 minutes at room temperature (RT). Cells were washed 3 times with 1X PBS, and blocking was performed with 3% (w/v) BSA and 3% (w/v) milk mixture (1:1) in PBST for 1 hour at RT. Further, cells were incubated with primary antibody of anti-ALFA (1:250) and anti-LAMP1 (1:250) in blocking buffer for overnight at 4 °C. The cells were then washed 3 times with PBST and counterstain with DAPI (1 uM) (#D1306, Invitrogen) for 15 minutes at RT, followed by 3 washes with 1X PBS and mounted on glass slide with ProLong Diamond (#36980, Invitrogen) anti-fading media. All images were captured by using Zeiss LSM900 confocal microscope with 63X objective (NA 1.4).

To perform the live cell staining of LRRC8A-ALFA and LysoTracker Red, LRRC8A-ALFA was transiently transfected into LRRC8A knockout C2C12 myoblasts. After 48 hours, live cells were pulsed with anti-ALFA-ATTO488 secondary antibody (1:100) for 5 minutes. Subsequently, the cells were replaced with fresh media and chased for 2 hours to allow the LRRC8A-ALFA-bound secondary antibody to undergo endocytosis. Further labelling with LysoTracker Red (#70083, Biotium, LysoView 550) was performed in the same cell culture media for 15 minutes. The cells were imaged by using STEDYCON Abberior microscope in a confocal mode with 100X objective. The fluorescent intensities of LRRC8A-ALFA and LysoTracker Red were measured using the line scanning tool in ImageJ.

### pH Measurement

Cells were grown on live cell dishes for LysoSensor Yellow/Blue DND-160 (#7545, Thermo Fisher Scientific) ratiometric imaging. For some experiments, myoblasts were differentiated into myotubes and freshly prepared Lysosensor (1 uM) was added to growth medium for 5 minutes (C2C12 cells) or 20 minutes (Primary muscle myotube) at 37°C. In a subset of experiments, differentiated myotubes were treated with Bafilomycin A1 (500 nM) for 4 hours prior to labelling with Lysosensor. To reduce background fluorescence, both labeling and imaging, were performed in phenol red-free DMEM. Lysosensor-stained cells were excited sequentially with 340 nm and 380 nm, and the resulting emissions were collected using a Fura-2 dichroic filter cube. The images were saved as TIFF files. The background was estimated using a rolling ball estimation method with a radius of 50 pixels in ImageJ. After background subtraction, the mean intensity of the entire field for the 340 nm and 380 nm images was estimated using the ‘Measure’ feature, and a ratio was calculated. In live cells, Lysosensor fluoresces brighter in the longer wavelength channel (380 nm) in lower pH (acidic) conditions, whereas at higher pH the emission shifts to shorter wavelengths (340 nm). Therefore, the 340/380 ratio is a good measure of pH of the organelle. Typically, 8-10 images were collected for each culture dish, and pH measurements were performed in triplicate for each condition. The 340 /380 ratio was plotted as a function of condition and one way ANOVA was used to estimate the significance of difference between different conditions.

### TEM Imaging and Lysosome Quantification

The TEM imaging and lysosome quantification were performed as described previously^44,45^. Briefly, cells were grown on coverslips, and before fixation, the cell culture media were removed. The cells were washed with a washing buffer (0.15 M cacodylate buffer with 2 mM CaCl₂) to remove any remaining cell culture media. Subsequently, cell fixation was performed with a fixative solution (2.5% glutaraldehyde, 2% paraformaldehyde, 0.15 M cacodylate buffer, 2 mM CaCl₂, pH 7.4) in an incubator for 15 seconds. The fixation was then continued overnight at room temperature on a shaker with gentle agitation. Samples were secondarily fixed in 1% osmium tetroxide and 1.5% potassium ferrocyanide in 0.1 M sodium cacodylate buffer for 30 minutes to 1 hour after an initial fixation and rinse. Following secondary fixation, samples were washed in 0.1 M sodium cacodylate buffer (pH 7.3) and diH2O, then incubated overnight at 4°C in 2.5% uranyl acetate. Dehydration was performed using an ethanol gradient, followed by infiltration with Eponate 12. Samples were cured at 70°C overnight, and sections were prepared using an ultramicrotome and counterstained with uranyl acetate and Reynold’s lead citrate. Transmission electron microscopy (TEM) images were acquired using either a JEOL JEM-1230 (120 kV) or a JEOL 1400 (80 kV) or Transmission Electron Microscope (JEOL JEM-1400 Plus, 120 kV). Lysosome area and circularity were quantified by ImageJ as described earlier^44,45^. Briefly, images were uploaded in TIFF format and divided into four quadrants using the ImageJ quadrant picking plugin. Two quadrants were randomly selected for detailed analysis. Three independent, blinded analysts quantified lysosomes and autophagosomes, averaging their results to minimize bias. A minimum of 7-10 cells were analyzed per sample to ensure reproducibility. Lysosome area and circularity were obtained by manual tracing using the freehand tool in NIH ImageJ.

### Lysosomal Immunoprecipitation

Lysosomal Immunoprecipitation from tissues or cells were adapted from the paper by Monther et al^46^. Briefly, differentiated myotubes were washed with cold 1X PBS, and scraped in 2 ml of homogenization buffer. The lysate was passed through a 26-gauge syringe 5-7 times and dounce homogenized with 6-8 strokes with a glass homogenizer under ice-cold conditions. The homogenized lysate was centrifuged at 6800 x g for 5 minutes at 4 °C, and the supernatant (PNS) fraction passed through 70 um pre-separation filter (#130-095-823, Miltenyi Biotech) equilibrated with homogenization buffer. The filtered PNS fraction was transferred to 50 uL of anti-HA (#88836, Thermo scientific) magnetic beads and rotated for 15-20 minutes in a cold room. The PNS-bound magnetic beads separated by using magnetic bead rack separator. The PNS-bound magnetic beads were washed 3 times with washing buffer. After washing, half of the PNS-bound beads were triton treated by using 60-70 ul lysis buffer. Subsequently, supernatant and beads were separated by magnetic separator, and further boiled with 4X Laemmli buffer.

To perform Lyso IP from mouse heart, the heat was dissected from the great vessels and atria under a dissecting microscope, and then washed with ice-cold PBS. The washed heart tissue was diced into small chunks using a razor blade and transferred to a cryotube with 500 µl of homogenization buffer [50 mM potassium chloride, 90 mM potassium gluconate, 1 mM EGTA, 50 mM Sucrose, 20 mM HEPES, 5 mM Glucose, pH 7.4, supplemented with protease and phosphatase inhibitors (Roche)] for further mechanical grinding using an IKA-T10 Basic Ultra-Turrax Disperser Homogenizer at maximum speed on ice. The lysate was transferred to a glass Dounce homogenizer to further homogenize the lysate with 20-30 strokes under ice-cold conditions. The homogenize lysate was transfer to a microcentrifuge tube, the volume was adjusted to 1 ml, and centrifuged at 6800 x g for 5 minutes at 4 °C. The supernatant or post-nuclear supernatant (PNS) was carefully transferred to a new tube and the volume was adjusted to 2 ml by adding homogenization buffer. The PNS fraction was passed through 70 um pre-separation filter (#130-095-823, Miltenyi Biotech) equilibrated with homogenization buffer. The filtered PNS fraction was then transferred to 50 uL of anti-Flag microbeads (#130-101-591, Miltenyi Biotech, µMACS™ and MultiMACS™ DYKDDDDK Isolation Kits) and rotated for 15 minutes in a cold room. The PNS-bound beads were then filtered through 20 um pre-separtion filter (#130-101-812, Miltenyi Biotech) pre-equilibrated with homogenization buffer. The filtered PNS-beads suspension was transferred to a MACS column on a MACS separator magnet and washed 3 times with 5 ml of wash buffer [1X PBS, 2 mM EDTA, 0.5% BSA, pH 7.4], allowing it to drain by gravity flow. After washing, the column was removed from the MACS separator, and 2 ml of homogenization buffer was added to elute the tagged protein bound to the beads. The eluted sample was subsequently centrifuged at 20,000 x g for 15 minutes to pellet the beads with bound lysosomes. Furthermore, to perform triton treatment, half of the beads were treated with 80 uL of lysis buffer [150 mM NaCl, 40 mM HEPES, 2.5 mM MgCl2, 1% TritonX-100, 2 mM EGTA, with proteinase/phosphatase inhibitors] for 15 minutes and centrifuged at 20, 000 x g for 15 minutes. The triton-treated supernatant, and pellet (beads) were then boiled with 4X Laemmli buffer for WB analysis.

### RNA Sequencing

Differentiated primary myotubes were solubilized in TRIzol and the total RNA was isolated using PureLink RNA kit (#12183018A, ThermoFisher Scientific) and column DNase digestion kit (#12185010, ThermoFisher Scientific). The RNA quality analysis and sequencing was performed by Genome Engineering & Stem Cell Center (GESC@MGI) at the Washington University in St. Louis. Total RNA integrity was determined using Agilent Bioanalyzer or 4200 Tapestation. Library preparation was performed with 10ng of total RNA with a Bioanalyzer RIN score greater than 8.0. ds-cDNA was prepared using the SMARTer Ultra Low RNA kit for Illumina Sequencing (Takara-Clontech) as per manufacturer’s protocol. cDNA was fragmented using a Covaris E220 sonicator using peak incident power 18, duty factor 20%, cycles per burst 50 for 120 seconds. cDNA was blunt ended, had an A base added to the 3’ ends, and then had Illumina sequencing adapters ligated to the ends. Ligated fragments were then amplified for 12-15 cycles using primers incorporating unique dual index tags. Fragments were sequenced on an Illumina NovaSeq X Plus using paired end reads extending 150 bases. Basecalls and demultiplexing were performed with Illumina’s DRAGEN and BCLconvert version 4.2.4 software. RNA-seq reads were then aligned to the Ensembl release 101 GRCm38 primary assembly with STAR version 2.7.9a. Gene counts were derived from the number of uniquely aligned unambiguous reads by Subread:featureCount version 2.0.3. Isoform expression of known Ensembl transcripts were quantified with Salmon version 1.5.2. Sequencing performance was assessed for the total number of aligned reads, total number of uniquely aligned reads, and features detected. The ribosomal fraction, known junction saturation, and read distribution over known gene models were quantified with RSeQC version 4.0.

All gene counts were then imported into the R/Bioconductor package EdgeR and TMM normalization size factors were calculated to adjust for samples for differences in library size. Ribosomal genes and genes not expressed in the smallest group size minus one samples greater than one count-per-million were excluded from further analysis. The TMM size factors and the matrix of counts were then imported into the R/Bioconductor package Limma. Weighted likelihoods based on the observed mean-variance relationship of every gene and sample were then calculated for all samples with the voomWithQualityWeights function and were fitted using a Limma generalized linear model with additional unknown latent effects as determined by surrogate variable analysis (SVA). The performance of all genes was assessed with plots of the residual standard deviation of every gene to their average log-count with a robustly fitted trend line of the residuals. Differential expression analysis was then performed to analyze for differences between conditions and the results were filtered for only those genes with Benjamini-Hochberg false-discovery rate adjusted p-values less than or equal to 0.05.

Sequencing results were uploaded and analyzed using BaseSpace (Illumina). Sequences were trimmed to 125bp using FastQ Toolkit (version 2.2.5). RNA-seq alignment (version 1.1.1) aligned the sequences against the *Mus Musculus* (UCSC mm10) genome. Transcript assembly and differential gene expression was determined by Cufflinks Assembly and DE (version 2.1.0). Ingenuity Pathway Analysis (QIAGEN) was used to analyzed significantly regulated genes using genes with >1.5-fold change in gene expression, and a false discovery rate of <0.05. Heat maps were generated using z-scores and plotted using GraphPad (version 10.2.1).

### Western blot

Cells were washed with ice cold 1X PBS and lysed in ice-cold RIPA lysis buffer (150 mM NaCl, 20 mM HEPES, 1% NP-40, 5mM EDTA, pH 7.5) with added proteinase/phosphatase inhibitor (Roche). The cell lysate was further sonicated 2-3 cycle (20% pulse frequency for 20 sec) and centrifuged at 13000 rpm for 20 min at 4°C. The supernatant was collected and estimated for protein concentration using DC protein assay kit (Bio-Rad). For immunoblotting, an appropriate volume of 4 x Laemmli (Bio-rad) sample loading buffer was added to the sample (10-20 μg of protein), then heated at 95°C for 5 min before loading onto 4-20% gel (Bio-Rad). Proteins were separated using running buffer (Bio-Rad) for 2 h at 110 V. Proteins were transferred to PVDF membrane (Bio-Rad) and membrane blocked in 5% (w/v) BSA or 5 % (w/v) milk in TBST buffer (0.2 M Tris, 1.37 M NaCl, 0.2% Tween-20, pH 7.4) at room temperature for 1 hour. Blots were incubated with primary antibodies (1:1000) at 4 °C overnight, followed by secondary antibody (Invitrogen, Goat-anti-Rat #31470, Goat-anti-mouse #170-5047, Rabbit-anti-mouse #D3V2A, Goat-anti-rabbit #170-6515, all used at 1:10000) at room temperature for one hour. Membranes were washed 3 times with TBST buffer and imaged by chemiluminescence (Pierce) by using a Chemidoc imaging system (BioRad). The images were further analyzed for band intensities using ImageJ software. β-Actin or GAPDH levels were quantified for equal protein loading.

### Metabolic Phenotyping

Mouse body composition (fat and lean mass) was measured by nuclear magnetic resonance (NMR); Echo-MRI 3-in-1 analyzer, EchoMRI, LLC) as described previously^20^. To measure glucose tolerance test (GTT), mice were fasted for 6 hours, and intraperitoneal injection of glucose (1g/kg body weight) administered. Glucose level was monitored from blood via tail bleeds using a glucometer (Bayer Healthcare LLC) at the indicated times. To measure insulin tolerance test (ITT), mice were fasted for 4 hours and after an intra-peritoneal injection of insulin (HumulinR, 1U/kg) glucose level was measured by glucometer at the indicated times.

### Hyperinsulinemic euglycemic glucose clamps

A sterile silicone catheter was introduced into the jugular vein under isoflurane anesthesia. Mice were allowed to fully recover from surgery for 5-6 days before undergoing hyperinsulinemic-euglycemic clamps. Animals showing impaired recovery (as evidenced by wound infection, weight loss over 10% compared to pre-surgery weight) were excluded from further experiments. Clamps were performed in 5 hours fasted, unrestrained, conscious mice by using a infusion swivel (#375/D/22QM, Instech, Plymouth Meeting PA) to allow free movement. Whole-body glucose flux was traced by infusion of D-[3-^3^H]-glucose. After an 80 min basal sampling period, insulin administration was initiated with a 32.5 mU/kg bolus infusion over 1 minute followed by 3.25 mU/kg/min continuous infusion. Simultaneously, 50% dextrose in saline was infused, adjusting the rate every 10 minutes to achieve a goal blood glucose of 150 mg/dL. Once the desired blood glucose level was stable (typically after 70-110 minutes), a glucose uptake tracer (^14^ C-2-deoxyglucose (2-DG) (Perkin Elmer, cat # NEC495001MC) was administered as a single bolus at a dose of 13 μCi over 1 minute at 75 minutes. ^14^C-2-deoxy-D-glucose-6-phosphate tracer enrichment was used to measure glucose uptake into specific tissues. Blood samples (∼20-25 μl) from tail-cut were collected at times 80, 90, 100, 110 and 120 mins, centrifuged, and plasma was transferred to a fresh tube and stored frozen at −80°C until use. After the clamp period, mice were anaesthetized with isoflurane, euthanized, tissues of interest were harvested, and frozen.

Glucose concentrations in the plasma and in the 50% dextrose infusates were measured using an enzymatic method (#GMD9, Analox Instruments Ltd, King William Street Amblecote Stourbridge). Plasma samples were treated with Ba(OH)_2_ and ZnSO_4_ and centrifuged at 16,000 g for 5 minutes. The supernatant was dried overnight at 50°C, then reconstituted in distilled water. Diluted ^3^H(3)-glucose infusate was processed in the same manner as plasma samples to serve as reference. Tracer (^3^H and ^14^C) activity was measured using liquid scintillation counter (Ultima Gold, Perkin Elmer, Waltham, MA). The obtained values were used for determination of the relevant glucose turnover parameters (Rd and Ra). Glucose appearance and disappearance rates were calculated using Steele’s equations^47^. Small (∼20-40 mg) tissue samples were treated with 0.5% perchloric acid, homogenized using a mechanical device (TissueLyzer, Qiagen, Hilden, Germany), then centrifuged at 4000g for 20 min. Supernatant was neutralized with KOH, then centrifuged at 10,000 g for 10 min. A portion of the resulting supernatant underwent liquid scintillation counting to determine total tracer (^14^C) activity (both phosphorylated and non-phosphorylated 2-DG). A separate portion was treated with Ba(OH)_2_ and ZnSO_4,_ then centrifuged at 16,000 for 5 min. The resulting supernatant underwent liquid scintillation counting to determine non-phosphorylated 2-DG activity (^14^C). The difference between radioactivity in first and second supernatant corresponds to the abundance of phosphorylated ^14^C 2-DG in the tissue which was used to determine tissue-specific glucose uptake by dividing by the integrated area of the plasma 2-DG activity curve. Plasma insulin concentration was measured using a chemiluminescence ELISA kit (#80-INSMR-CH01, Stellux, from ALPCO).

#### Statistics

Data are represented as mean ± s.e.m. Two-tail paired or unpaired Student’s t-tests were used for comparison between two groups. For three or more groups, data were analyzed by one-way ANOVA and Tukey’s *post hoc* test. For GTTs and ITTs, 2-way analysis of variance (Anova) was used. A p-value < 0.05 was considered statistically significant. *, ** and *** represents a p-value less than 0.05, 0.01 and 0.001 respectively.

## Author contributions

Conceptualization: R.S., A.K.

Formal Analysis: A.K., L.X., R.C., Y.Z., N.A., J.D.T., J.H., E.F., D.R., K.M.H., H.L., A.H.J.

Investigation: A.K., L.X., R.C., Y.Z., N.A., J.D.T., J.H., E.F., D.R., K.M.H., H.L., A.H.J., M.H., H.X.

Writing – Original Draft: A.K., R.S.

Writing – Review & Editing: R.S., A.K., C.E.G., G.M., A.D., A.W.N., A.H.J., E.D.A.

Funding Acquisition: R.S., C.E.G., G.M., A.D., E.D.A.

Visualization: A.K., C.E.G., A.H.J., R.S.

Resources: R.S., G.M., E.D.A., D.A.

Supervision: R.S., A.D., G.M.

## Acknowledgements

We thank the Fraternal Order of Eagles Diabetes Research Center at the University of Iowa for performing euglycemic hyperinsulinemic clamps. We gratefully acknowledge Gregory Strout, and John Wullf II for their assistance in electron microscopy studies conducted at the Washington University Center for Cellular Imaging (WUCCI), which is supported in part by Washington University School of Medicine, The Children’s Discovery Institute of Washington University and St. Louis Children’s Hospital (CDI-CORE-2015-505 and CDI-CORE-2019-813), the Foundation for Barnes-Jewish Hospital (3770) and the Washington University Diabetes Research Center (NIH P30 DK020579). We thank the Genome Engineering & Stem Cell Center (GESC@MGI) at the Washington University in St. Louis for reagent validation services. We thank University of Iowa electron microscopy core for TEM imaging assistance. We also thank Dr. Thomas Jenstch for kindly sharing LRRC8B, LRRC8D, and LRRC8E rabbit polyclonal antibodies. We thank Antony Lurie and Dr. Stephen G. Brohawn for kindly providing the LRRC8A-ALFA-IRES-EGFP and LRRC8A-ALFA plasmids. This work was supported by NIH NIDDK R01DK106009 (R.S.), R01DK126068 (R.S.), R01DK127080 (R.S.), R01-DK115791 (AWN), NIH NHLBI R01HL168600 (R.S.), R01HL107594 (A.D.), R01HL168600 (A.D.), R01HL108379 (E.D.A), K08HL163469 (D.R.). Department of Veteran Affairs I01 BX005072 (R.S.), I01BX004235 (A.D), I01BX005065 (A.D.), I01BX005981 (A.D.). CZI Science Diversity Leadership grant (2022-253529, A.H.J).

## Competing interests

R.S. is co-founder of Senseion Therapeutics, Inc., a start-up company developing SWELL1 modulators for human disease. The remaining authors declare no competing interests.

## Supplementary figures

**Supplementary Figure 1.**
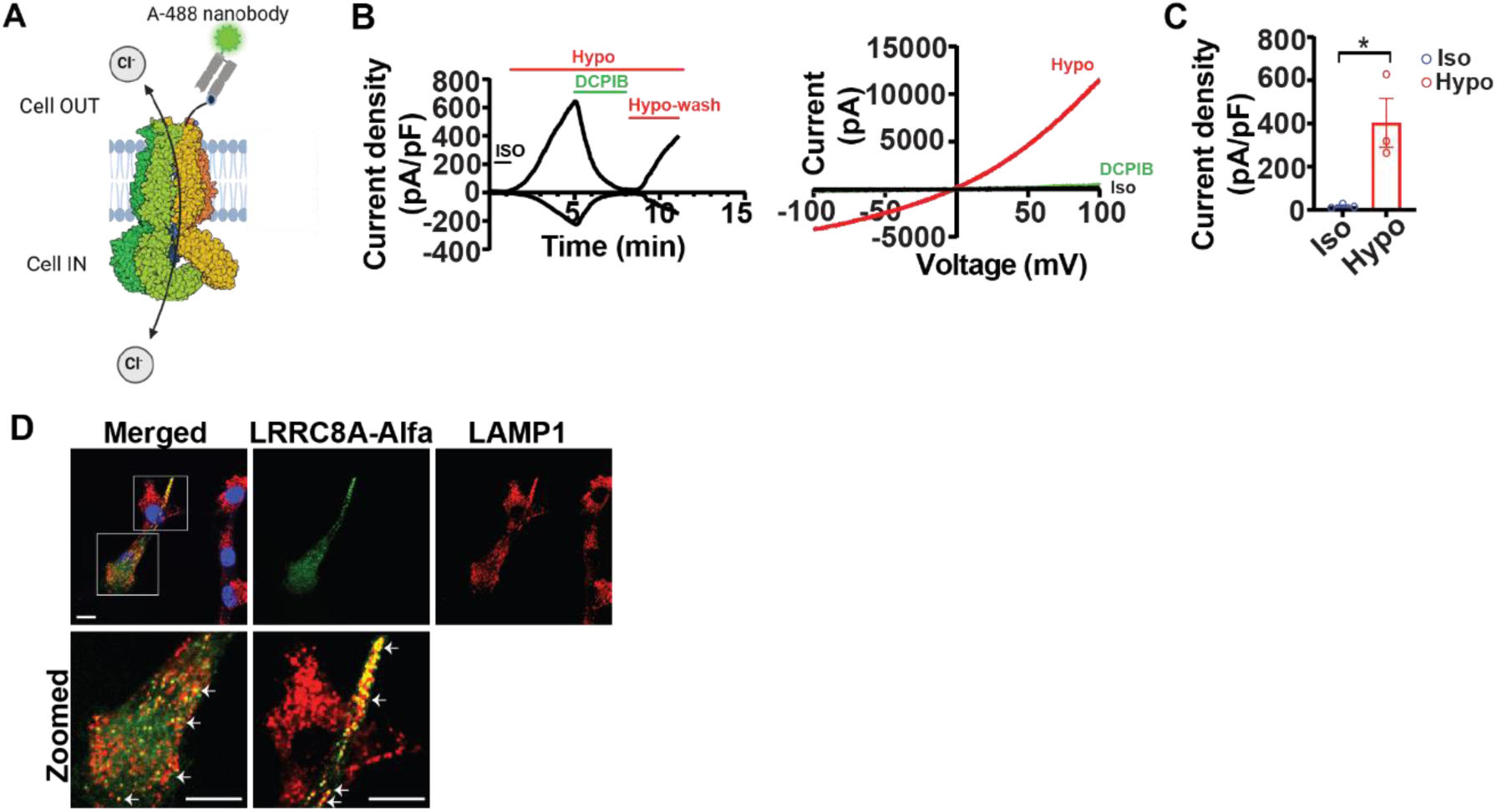
Functional characterization of LRRC8A-alfa tagged protein in cells A, Cartoon depicting a heterohexameric structure of LRRC8A complex showing alfa epitope (13 aa) were inserted on first extracellular loop of mouse LRRC8A protein by replacing 77-90 aa. **B & C,** LRRC8A-alfa-IRES-EGFP and LRRC8C-P2A-mCherry were co-expressed transiently in 5 KO (LRRC8A/B/C/D/E KO) HeLa cells, and whole cell Current-voltage were measured. A voltage-ramp −100 to +100 mV were applied in presence/absence of isotonic (300 mOsm) and hypotonic (210 mOsm) solution and subsequent inhibition of VRAC current by 10 uM of DCPIB. **D,** LRRC8A-alfa were transiently transfected in LRRC8A KO C2C12 myoblast cells, and immunostaining was performed with anti-LAMP1 and anti-alfa nanobody. Confocal imaging shows colocalization of lysosomal protein LAMP1 (Red) with LRRC8A-alfa (Green) (white arrow). Scale bar: 10 µm.

**Supplementary Figure 2.**
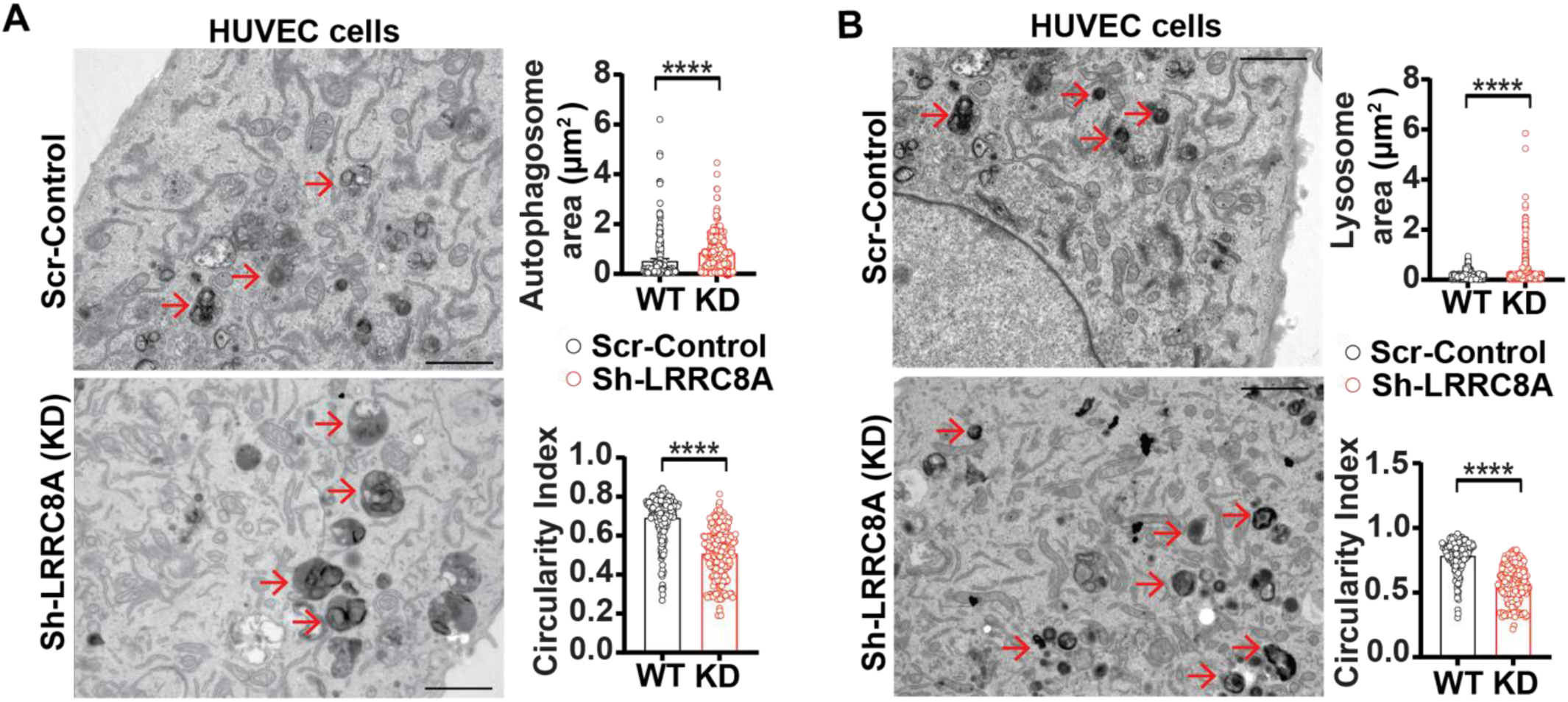
LRRC8A depleted cells show enlarged lysosomes A&B, TEM images of WT human umbilical vein endothelial cell (HUVEC) and LRRC8A KD HUVEC showing enlarged autophagosome and lysosomes. The autophagosome area (WT=401, LRRC8A KD=586 autophagosomes), lysosome area (WT=497, LRRC8A KD=556 lysosomes) and circularity index for autophagosome (WT=419, LRRC8A KD=572 autophagosomes) and lysosomes (WT=254, LRRC8A KD=213 lysosomes) shown on the right side of images. Scale bar: 2 µm.

**Supplementary Figure 3.**
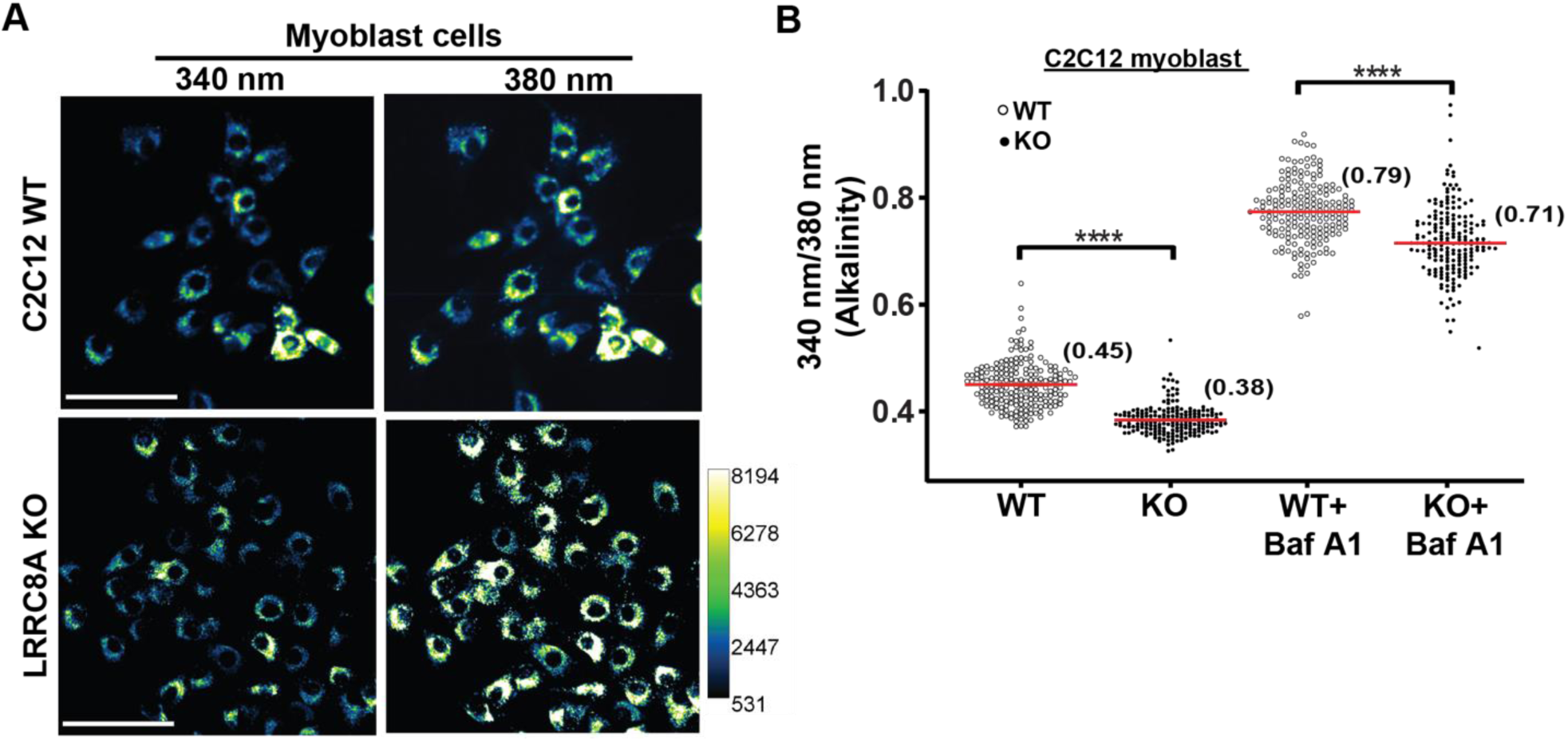
LRRC8A depletion decreases lysosomal pH A, Fluorescence images of Lysosensor labeled images of WT and LRRC8A KO myoblasts. Scale bar: 100 µm. **B** Ratiometric (Ex340/Ex380) intensity quantification of Lysosensor stained images from 5-6 different fields of view (n=3 independent experiment).

**Supplementary Figure 4.**
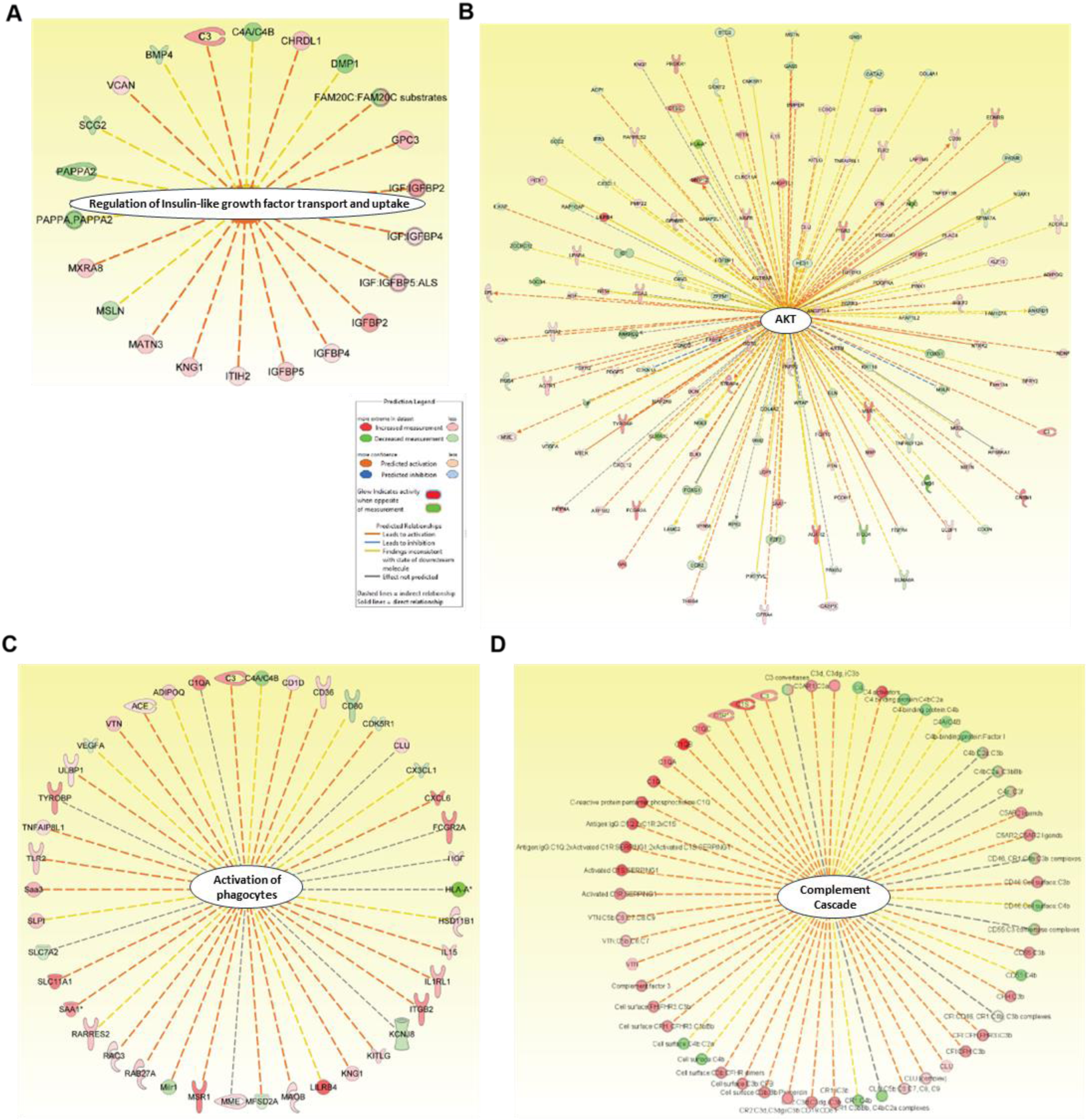
RNA transcriptome analysis of lyso-null LRRC8A (LL:AA) shows altered cellular signaling pathway A-D. Interaction networks identified affected cellular signaling pathway in LL:AA such as regulation of (A) Insulin-like growth factor transport and uptake, (B) AKT signaling, (C) activation of phagocytosis, (D) complements cascade. Network associated gene legends and symbol indicated by predicted legend figure.

**Supplementary Figure 5.**
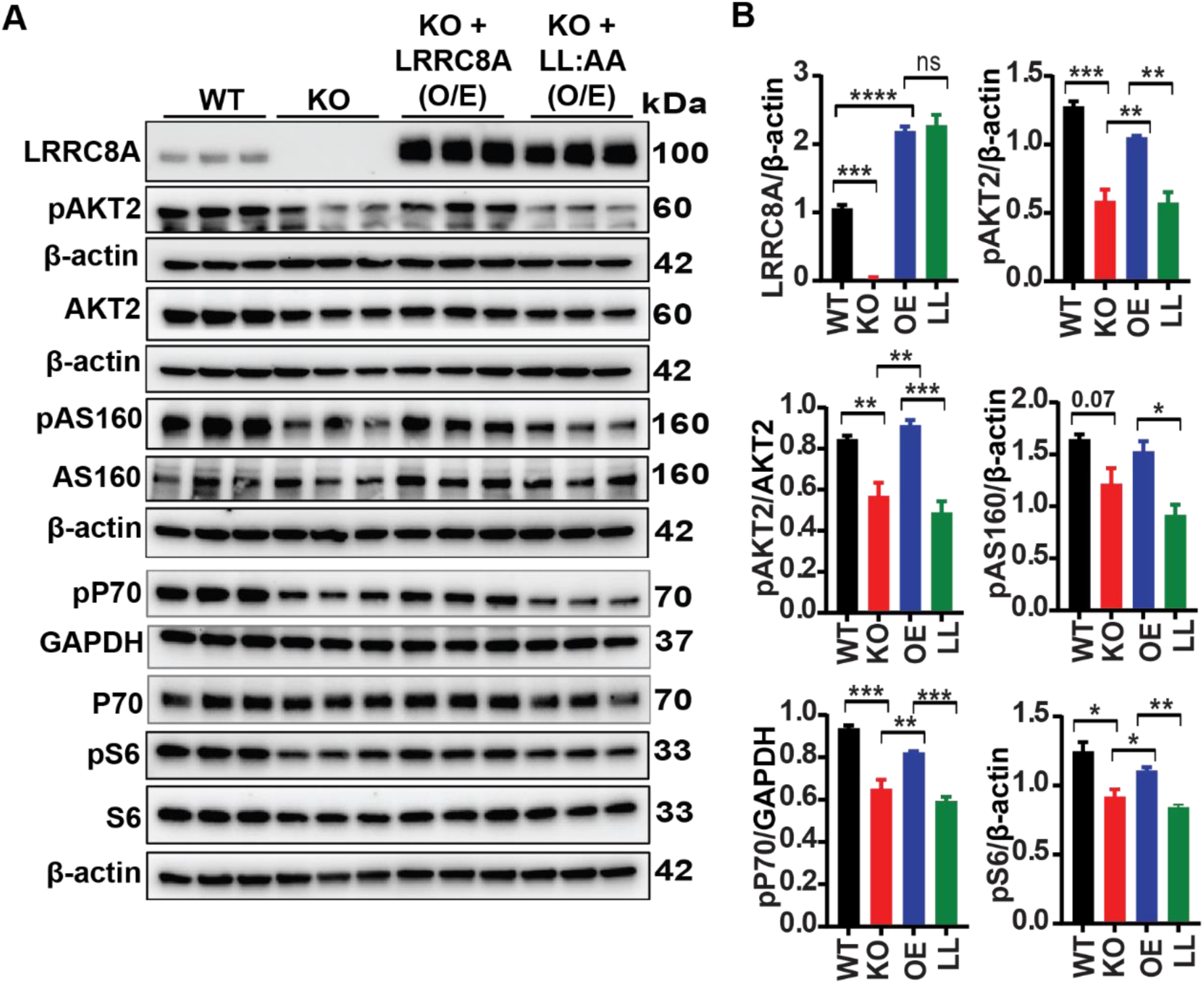
Re-expression of LRRC8A-LL:AA in LRRC8A null cells fail to restore PI3K-AKT-mTOR signaling A, Western blots of LRRC8A, pAKT2, AKT2, pAS160, AS160, pP70, P70, pS6, S6, β-actin and GAPDH in WT C2C12, LRRC8A KO C2C12, LRRC8A KO + LRRC8A-3flag, LRRC8A KO + LL:AA-3flag C2C12 myotubes at basal condition. **B,** Densitometry quantification of WB (A). Statistical significance between the indicated group were calculated with one-way ANOVA, Tukey’s multiple comparisons test. Error bars represent mean ± s.e.m. *, p<0.05, **, p<0.01, ***, p<0.001, ****, p<0.0001. n = 3, independent experiments.

